# INF2-mediated actin filament reorganization confers intrinsic resilience to neuronal ischemic injury

**DOI:** 10.1101/2021.10.01.462790

**Authors:** Barbara Calabrese, Steven Jones, Yoko Yamaguchi-Shiraishi, Michael Lingelbach, Uri Manor, Tatyana M Svitkina, Henry N Higgs, Andy Y Shih, Shelley Halpain

## Abstract

During early stages of ischemic brain injury, glutamate receptor hyperactivation mediates neuronal death via osmotic cell swelling. Here we show that ischemia and excess NMDA receptor activation – conditions that trigger neuronal swelling -- cause actin filaments to undergo a rapid and extensive reorganization within the somatodendritic compartment. Normally, F-actin is concentrated within dendritic spines, with relatively little F-actin in the dendrite shaft. However, beginning <5 min after incubation of neurons with NMDA, F-actin depolymerizes within dendritic spines and polymerizes into long, stable filament bundles within the dendrite shaft and soma. A similar “actinification” of the somatodendritic compartment occurs after oxygen/glucose deprivation *in vitro*, and in mouse brain after photothrombotic stroke *in vivo*. Following transient, sub-lethal NMDA exposure these actin changes spontaneously reverse within 1-2 hours. A combination of Na^+^, Cl^-^, water, and Ca^2+^ entry are all necessary, but not individually sufficient, for induction of actinification. Spine F-actin depolymerization is also required. Actinification is driven by activation of the F-actin polymerization factor inverted formin-2 (INF2). Silencing of INF2 renders neurons more vulnerable to NMDA-induced membrane leakage and cell death, and formin inhibition markedly increases ischemic infarct severity *in vivo*. These results show that ischemia-induced actin filament reorganization within the dendritic compartment is an intrinsic pro-survival response that protects neurons from death induced by swelling.

## INTRODUCTION

Ischemic stroke results from occlusion of cerebral blood vessels, causing brain tissue injury due to loss of oxygen and glucose supply ^1^. It is a leading cause of death and chronic disability, and has enormous public health implications, especially recently in the context of Covid-19, which significantly increases a severely ill patient’s risk of stroke ^2–5^. Beyond anticoagulant therapies, there are relatively few emergency treatment options for stroke. This highlights the need for a deeper understanding of the biological cascades ensuing from an ischemic event, including cellular pro-survival mechanisms that could minimize the ultimate brain tissue damage.

Although apoptosis accounts for much of the delayed neuronal death that occurs over days and weeks following a stroke, most of the neuronal death that occurs in the early hours after a stroke is due to a pathological swelling of neurons that leads to disruption of plasma membrane integrity ^6–9^. Neuronal swelling, which is also called cytotoxic edema, is triggered when catastrophic ATP depletion perturbs ionic balance, leading to a massive influx of ions through multiple entry routes, with cation entry through NMDA receptors and chloride entry through the SLC26A11 ion exchanger playing major roles ^10^. Neuronal depolarization spreads in waves from the site of initial ischemia via local release of glutamate, thereby exacerbating and extending the initial damage^11^. NMDA receptor hyperactivation plays an especially critical role, as administration of NMDA receptor antagonists before or even just after the onset of ischemia results in significantly reduced infarct volume in experimental models^11–13^.

Neurons, like most other cells, respond to cellular stress and injury, including ischemic injury, by mounting a series of pro-survival responses, with changes in organelles, reduction in protein synthesis, and activation of pro-survival genes^14^. In contrast, the pro-survival functions of the cytoskeleton are less well characterized. Here we describe an extensive reorganization of neuronal filamentous actin (F-actin) that occurs in response to stroke, oxygen and glucose deprivation, or NMDA receptor hyperactivation. F-actin is rapidly depolymerized within dendritic spines^14–16^. Simultaneously, we show here, it polymerizes extensively within the soma and dendrites. This result was surprising given the usual ATP dependence of actin polymerization and the reduced ATP availability during ischemia. We find that this F-actin response is pro-survival and selectively triggered by conditions that elicit neuronal cytotoxic edema. The F-actin build-up in the soma and dendrites results in long, slowly turning over actin filaments that persist while the stress is present. However, the F-actin reorganization spontaneously reverses if the stress is transient. We demonstrate that activation of inverted formin-2 (INF2) is a key mediator of this neuronal pro-survival response.

## RESULTS

### Actin reorganization is induced by oxygen & glucose deprivation *in vitro*

To investigate actin filament responses to stroke-like conditions we exposed cultured rat brain hippocampal neurons to oxygen and glucose deprivation (OGD). OGD is a well-established *in vitro* model for investigating neuronal cellular and subcellular responses to hypoxia and ischemia ^17^. OGD induced a dramatic reorganization of F-actin within the soma and dendrites of neurons. Over 2-6 hours an increasing fraction of neurons showed a substantial loss of phalloidin staining for F-actin within dendritic spines, where F-actin is normally concentrated, and an aberrant accumulation of F-actin within the somatodendritic compartment (Fig 1A-C). Filaments accumulated throughout the interior of the soma and proximal dendrites, and to various extents within more distal dendrites.

**Figure 1.**
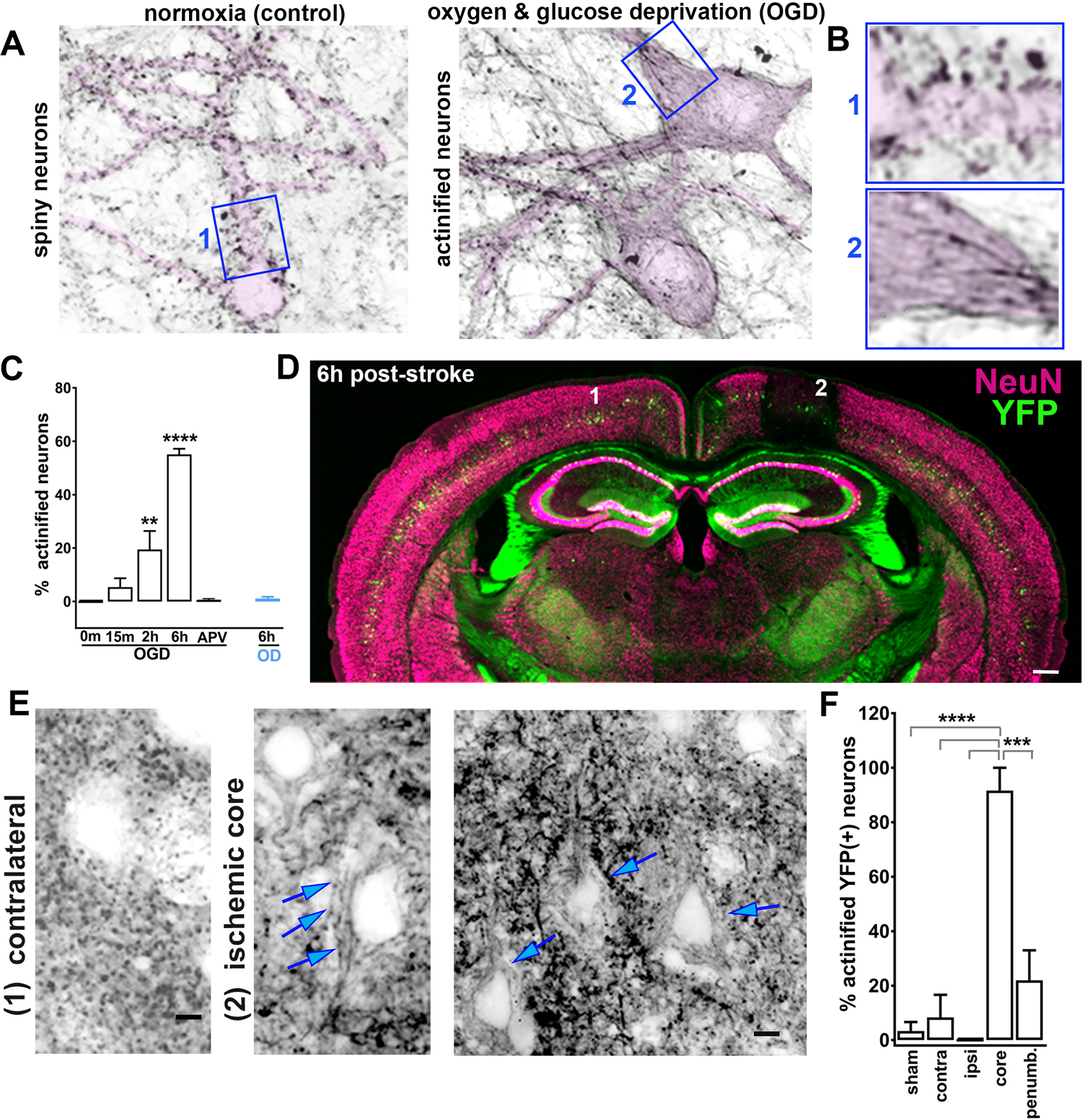
Ischemia-induced actin filament reorganization. A,B. <u>F-actin reorganization induced by oxygen and glucose deprivation (OGD)</u>. Cultured rat hippocampal neurons were stained using Alexa Fluor 647-phalloidin following incubation in the absence or presence of 6h oxygen and glucose deprivation (OGD), a standard *in vitro* model for ischemia (**A**). Fluorescence image is inverted for optimized display; the somatodendritic region of selected individual neurons in each panel are highlighted in *pink* to emphasize dendritic morphology (see details in Materials and Methods). Note that in the control condition F-actin is concentrated mainly in dendritic spines (dark puncta) along the dendrites, while OGD induces a decrease in spine F-actin and an accumulation of linear bundles of F-actin within the soma and dendrite of many neurons (“actinification”). Scale bar, 5 µm. (**B**) Numbered insets, corresponding to the blue boxed regions in **A**, are enlarged to show the different somatodendritic organization of the F-actin in the two respective experimental conditions, without the interference of processes from neighboring neurons (actinified or spiny). Width of blue boxes = 17 µm. (**C**) Quantification of the fraction of actinified neurons in control vs. OGD or OD (oxygen deprivation alone) for the indicated durations; the NMDA receptor antagonist (2R)- amino-5-phosphonovaleric acid (APV) prevented actinification seen with 6 hr OGD. Data are represented as mean ± SEM; n= 4 coverslips per condition. ** p<0.01;****p<0.0001; one-way ANOVA, Dunnett’s multiple comparison *post-hoc* test. **D-F.** <u>F-actin reorganization induced by photothrombotic stroke; mouse brains fixed and stained 6h post stroke induction</u>. (**D**) Coronal brain section of a Thy1-GFP (green) expressing mouse stained for the neuronal cell body marker NeuN (*magenta*). Cortical regions marked “1” and “2” indicate the areas from which the corresponding enlarged images in **E**, below are shown. Note: the infarct region is readily identified as the area of significant loss in NeuN staining, although the cell bodies of Thy1-GFP-positive neurons are still detectable (see also Suppl. Fig. S2). Scale bar, 350 µm. (**E**) Alexa-Fluor-647-Phalloidin staining of individual neurons within the somatosensory cortex within (2) the infarcted region (two panels shown at different magnification, *middle* and *right*, or (1) within the corresponding control region in the hemisphere contralateral to the stroke (single panel on the *left*). *Blue arrows* in the *middle* and *right* panels indicate the accumulation of linear actin bundles within the somatodendritic region of YFP-positive layer 5/6 neurons within the ischemic zone. In contrast, the F-actin staining within the control, contralateral region (*left*) exhibits the expected punctate pattern, where each punctum corresponds to a dendritic spine (see also Suppl. Fig. S3, demonstrating the correspondence of F-actin puncta to dendritic spines). Scale bars, 5 µm (*left* and *middle)*, 8 µm (*right)*. (**F**) Quantification of the fraction of YFP-positive layer 5/6 neurons exhibiting actinification under the following conditions: sham-operated mice (somatosensory cortex); stroke-induced mice, within the corresponding contralateral cortex; stroke-induced mice within temporal cortex ipsilateral to the infarct region; stroke-induced mice, within the ischemic core of the infarct; stroke-induced mice within the penumbral region of the infarct (as defined in Materials and Methods). Data are represented as mean ± SEM; n=3 animals each, sham vs. stroke; *** p<0.001;****p<0.0001, one-way ANOVA, Tukey’s multiple comparison *post-hoc* test.

Because in most spiny neurons the soma and shaft of the dendrite is fairly devoid of F-actin relative to dendritic spines, we call this novel phenomenon the “actinification” of the neuron. OGD-induced somatodendritic actinification was completely blocked in the presence of the NMDA receptor antagonist (2R)-amino-5-phosphonovaleric acid (APV; Fig 1C). Interestingly, oxygen-deprivation alone was insufficient to induce actinification within the same time frame (Fig 1C), perhaps because neurons sustain partial energy production via glycolysis for extended periods ^18^. Together, these results suggest that actinification is induced by a catastrophic loss of ATP, leading to excess glutamate release and NMDA receptor hyperactivation.

### Actin reorganization induced after micro-infarct (stroke) in vivo

We investigated whether ischemia *in vivo* also induces neuronal actinification using either wildtype mice or transgenic Thy1 promoter-driven YFP expressing mice, the latter of which allowed us to readily identify dendritic arbor morphology in layer 2/3 and 5/6 cortical pyramidal neurons. Photothrombotic occlusion in single penetrating arterioles within the somatosensory cortex of mouse or rat brain induces small infarcts with well-defined borders^12, 19^. Such strokes are greatly attenuated by application of NMDA antagonists before or immediately following blood vessel occlusion ^12^. Within 2-6 hours the infarct region induced by unilateral single vessel photothrombosis was identifiable using a variety of markers, including fluorescent hypoxyprobe to detect severely hypoxic tissue, anti-IgG to detect vascular leakage, and reduced immunostaining for the neuron-specific proteins MAP2 and NeuN ^20, 21^ (Fig 1D; Suppl Fig S1). We noted that, consistent with a previous report ^21^, the strongly reduced immunoreactivities for NeuN at this early time point reflected a loss of antibody staining, i.e., epitope loss, rather than a reduction in overall cell number, since DAPI staining for nuclear DNA and cytoplasmic YFP labeling were retained in neurons lacking NeuN (Suppl Fig S2). Actinification was detected within infarcts induced in either wildtype or Thy1-YFP mice.

Phalloidin staining revealed that in control brain tissue, as in control cultured neurons, the majority of F-actin was concentrated in dendritic spines (Fig 1E, Suppl Fig S3). By 4-6 hours after arteriole occlusion, the dendrites of GFP-expressing layer 5/6 pyramidal neurons showed a substantial loss or shrinkage of dendritic spines and the appearance of dystrophic dendrites, (Suppl Fig S4). Low-magnification imaging of the infarct region indicated there was a time-dependent decrease in overall F-actin concentration, consistent with a loss of dendritic spine F-actin (Suppl Fig S5). However, high magnification imaging of individual layer 5/6 YFP-positive neurons showed aberrant accumulations of F-actin bundles (Fig 1E). Quantitative analysis revealed that within the ischemic core of the infarct (defined by the region of NeuN loss) there was a significant increase in the number of actinified neurons compared to control neurons in either sham treated mouse brains, or in the stroke condition within contralateral cortex, or within ipsilateral temporal lobe cortex distant from the infarct region. Within the core of the infarct, nearly 90% of the YFP-labeled neurons showed actinification. (Fig 1F).

Dendritic spine shrinkage and spine F-actin loss have been described previously as an early response to stroke or to NMDA receptor hyperactivation,^15, 16, 22, 23^ and mechanisms contributing to spine F-actin loss following strong NMDA receptor activation have been investigated previously^16, 24–26^. However, the significant accumulation of F-actin in the somatodendritic compartment was unexpected, given that F-actin polymerization typically involves ATP consumption, and would logically seem to be disfavored in conditions of hypoxic cellular stress, when energy supplies become severely limited. We therefore focused on understanding the mechanism and relevance of somatodendritic actinification.

### NMDA receptor hyperactivation induces actinification within minutes

Since an NMDA receptor antagonist completely prevented somatodendritic actinification following OGD, we asked whether actinification could be triggered via direct activation of NMDA receptors. Incubation of cultured hippocampal neurons with 50 µM NMDA induced a time-dependent increase in neuronal actinification, accompanied by extensive loss of dendritic spine F-actin and shrinkage of spines, similar to that seen following OGD *in vitro* and stroke *in vivo* (Fig 2A). However, the time course for the actinification response was greatly accelerated. Based on a time series collected using fixed cultures, half-maximal actinification (the time at which 50% of all neurons were actinified) occurred after 5-10 minutes (average t_1/2_ = 7 min) in the sustained presence of NMDA, reaching nearly 100% by 60 minutes (Fig 2B).

**Figure 2.**
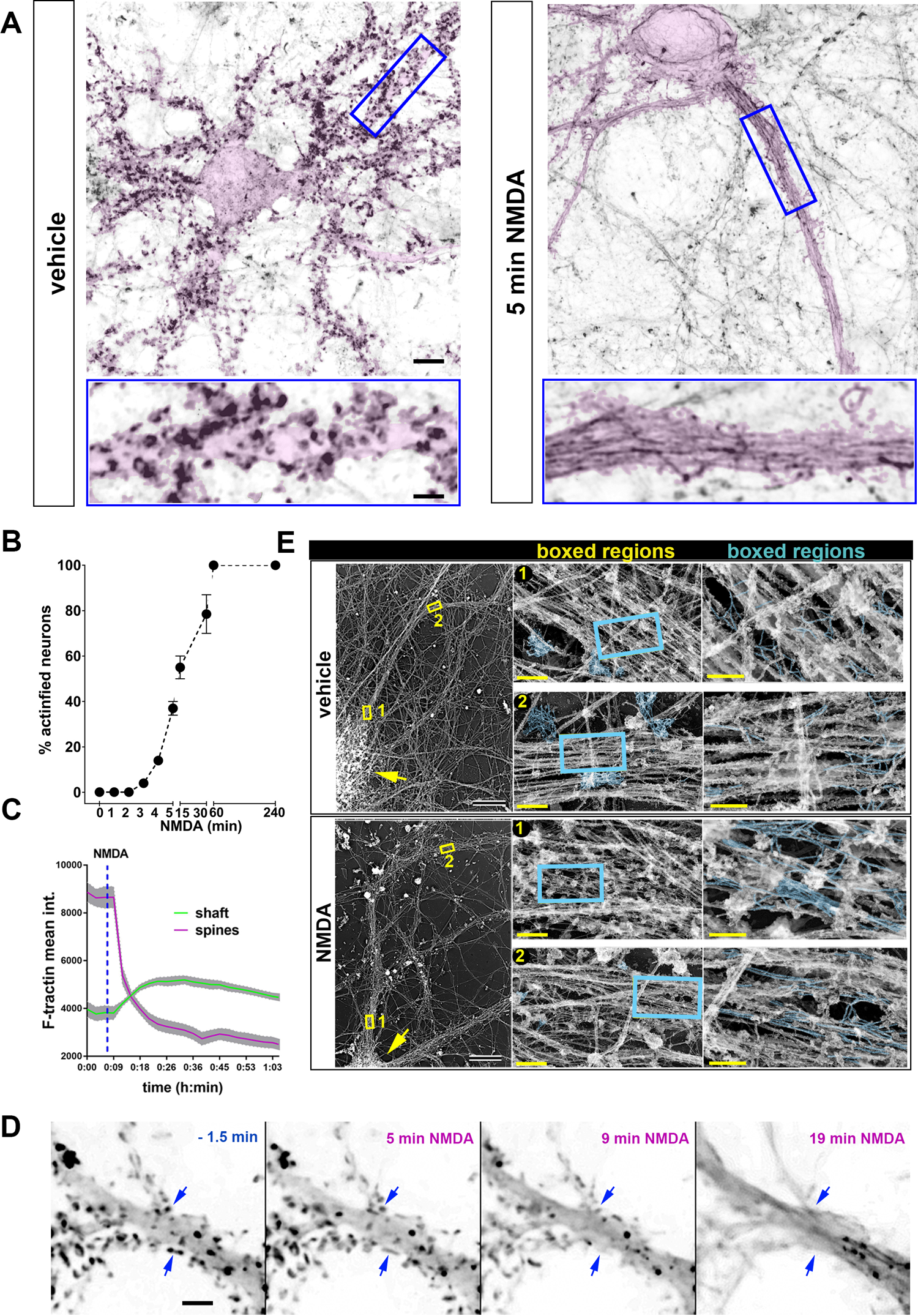
Rapid actinification of cultured hippocampal neurons induced by sub-lethal hyperactivation of NMDA receptors: time course and ultrastructure. Incubation of hippocampal cultures induces actinification within 5 min. Selected images of hippocampal neurons transfected with membrane-tagged GFP (*pink highlight*), to emphasize the dendritic morphology, and stained with Alexa Fluor 647-phalloidin to monitor F-actin localization (*reverse grayscale*). *Left*: Control culture fixed after 5 min incubation with vehicle alone; *Right*: stimulated culture fixed after 5 min incubation with 50 μM NMDA. Insets below show enlarged regions boxed in the upper portion of the figure. Scale bars, 8 µm (above); 2.8 µm (below). (**B**) Time course of actinification, expressed as percentage of all neurons that display the characteristic accumulation of actin bundles within the soma and dendrite. Data are mean ± SEM; n=4 replicates for 0 and 5 min; n=2 replicates for all the other time points. (**C**) Live imaging of individual hippocampal neurons hows that F-actin loss in the spines occurs in parallel with F-actin accumulation in the adjacent dendrite shaft. Time course from a representative neuron where mApple-F-tractin intensity was monitored over time in the dendrite shaft (*green line*) vs. the adjacent spines (*magenta line*) during NMDA-induced actinification. Plotted lines represent mean ± SEM from region of interest (ROIs) taken across different regions of the same dendrite and different spines of a single neuron. (**D**) A time-lapse montage of a dendritic region expressing Lifeact-mRFP at selected time points before and after exposure to 50 µM NMDA. Scale bar: 5 µm. *Blue arrows* point to examples of spine F-actin decreasing around the same time as the increase in F-actin signal in the shaft. (**E**) Platinum replica electron microscopy (PREM) demonstrates that actinification induces accumulation of linear, unbranched actin filaments within the dendrite. Hippocampal cultures were incubated for30 min in the vehicle or 50 µM NMDA as indicated, prior to fixation and processing for PREM, as described in Materials and Methods. A neuron representative of each condition is shown at *left*; yellow arrows indicates the position of the cell soma; yellow boxes correspond to two regions of the dendrite that are displayed at 70x higher magnification in the *middle* panels, and at a further 3x higher magnification in the *right* panels. Actin filaments identified by their thickness are highlighted in *cyan*. In control neurons (vehicle) numerous clusters of short, moderately branched actin filaments are observed in presumptive dendritic spines; linear actin filaments are occasionally detectable within the dendrite shaft. In contrast, in NMDA-treated neurons there is a substantial decrease in the frequency of spine-like protrusions containing branched actin filament networks, and numerous long, unbranched filaments were detected within the dendritic shaft of NMDA-treated cultures (*cyan* labeled structures within *cyan* boxed regions). Scale bars, 20 µM for left panels, 500 nm for middle panels, and 200 nm for right panels.

### Time-lapse imaging of actinification

We carried out time-lapse imaging of individual hippocampal pyramidal neurons using either Lifeact-mRFP, mApple-F-tractin, or the small molecule dye far-red silicon-rhodamine actin. The time course of live neurons undergoing actinification was consistent with the time course determined using fixed populations of neurons. Dendrite actinification occurred in a time frame that closely overlapped with the observed decrease in dendritic spine F-actin (Fig. 2C, D). As observed in fixed samples, live imaging showed that neurons underwent actinification after variable delays ranging from 3-20 minutes post-NMDA. However, once detectably initiated, actinification proceeded rapidly and with a similar time course.

### Ultrastructural analysis indicates that actinification results in long, unbranched actin filaments

The rapid time course of actinification initiated by NMDA suggests that the aberrant accumulation of F-actin filaments is likely due to an enzymatically-driven process, rather than the slower aggregation type process that governs build-up of pathological aggregates seen in neurodegenerative diseases such as Alzheimer’s. To investigate the characteristics of actinified dendrites at the ultrastructural level, we carried out platinum replica electron microscopy (PREM) on control cultures and compared them to cultures incubated with 50 µM NMDA for 5 or 30 min (Fig 2E). PREM images from control neurons revealed numerous clusters of short, moderately branched actin filaments localized within dendritic spine-like structures along the dendrite shaft, consistent with previous studies ^27^. Within the dendrite shaft of control neurons, actin filaments were relatively sparse. Long filaments running parallel to the main dendrite axis were rarely observed, and when detected they usually exhibited occasional branches. In contrast, and in agreement with our light microscopy observations, PREM images of dendrite shafts from NMDA-treated cultures revealed numerous instances of long, mostly unbranched filaments, which often formed irregular bundles. In parallel, there was a substantial decrease in the frequency of spine-like protrusions containing branched actin filaments that were common in control neurons. These observations suggest that dendrite actinification is driven by a process that disassembles branched actin filaments in spines and polymerizes F-actin into unbranched, rather than branched, filaments in dendrite shafts.

### Actin filaments induced by NMDA are extremely stable

Once formed, actin filaments in the actinified neuronal somata and dendrites appeared to be highly stable. We tested whether incubation with latrunculin A would accelerate the removal of the actinified F-actin in the dendrite shaft. Actin filaments within dendritic spines turnover with a half-time of < 1 minute ^28, 29^. We reasoned that if a similarly high turnover rate of F-actin characterizes the actinified dendritic compartment, we should observe that application of the G-actin sequestering compound Latrunculin A – applied <u>after</u> actinification has occurred -- would greatly speed the net disassembly of the filaments. However, our observations did not support this hypothesis. Incubation with 2 µM LatA for up to 2 hours after actinification induction caused no detectable decrease in the percentages of actinified neurons, nor loss of F-actin staining intensity within individual neurons in the soma and dendrites (Fig 3 A-C). Note that we first confirmed that LatA would prevent actinification, as expected, when applied before actinification was induced by NMDA (Fig. 3A), consistent with actinification being an actin monomer-dependent polymerization process. The resistance of filaments to latrunculin added post-actinification suggests that the newly polymerized F-actin induced by NMDA is extremely stable in the continued presence of NMDA.

**Figure 3.**
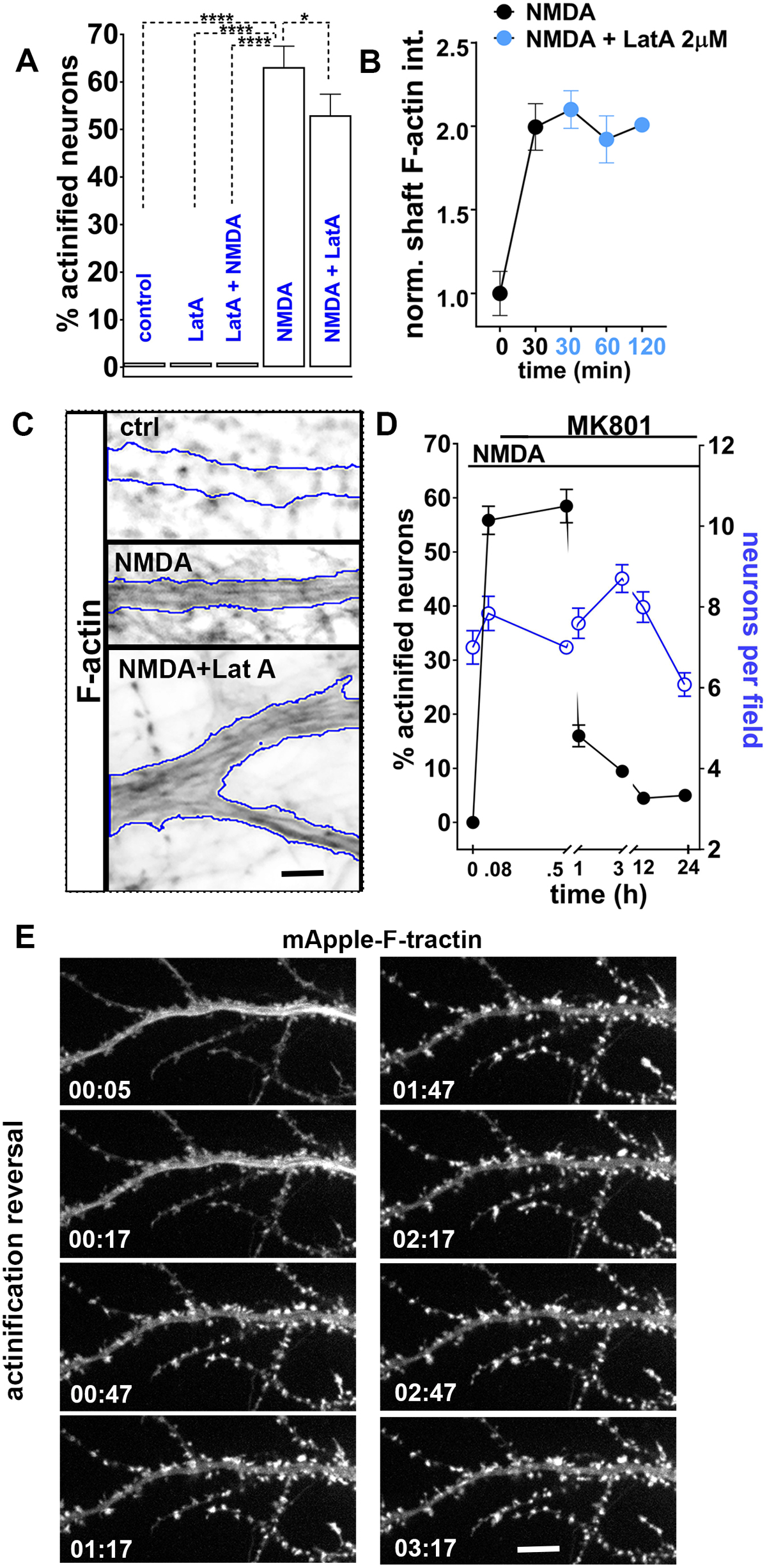
Actinification results in highly stable actin filaments that spontaneously disassemble upon NMDA stress removal. (**A**) Sequestration of actin monomer by latrunculin A (LatA; 2 µM) prevents actinification when applied 10 min before adding NMDA (“LatA + NMDA”), but does not reverse actinification when added for up to 2 hr following incubation in 50 µM NMDA for 5 min (“NMDA + LatA”). Results were quantified as percentage of actinified neurons; data represented as mean ±SEM; *p<0.05; ****p<0.0001; one-way ANOVA, followed by Tukey’s multiple comparison *post-hoc* test; 2 replicates per condition. (**B**) Quantification of the phalloidin staining intensity within the proximal dendrite of neurons exposed to 50 µM NMDA for 30 min followed by 2 µM LatA for the indicated times. Data are mean ±SEM; n = 20 neurons per time point. (**C**) Selected dendritic regions stained for phalloidin illustrating that NMDA-induced actinification persists after incubation with LatA for 2 hr (borders of the dendrite are outlined in *yellow*). Scale bar, 4 µm. (**D**) Reversibility of NMDA-induced actinification. Results were quantified as the fraction of actinified neurons (*black circles*) in the continued presence of 50 µM NMDA for the indicated times; the NMDA antagonist MK-801 (25 µM) was added after incubation with NMDA for 5 min to block further NMDA receptor activation, and cultures were fixed after the indicated times. In the same samples the total number of neurons per field (*blue circles*) were quantified as an indicator of neuronal viability, to rule out that the decrease in actinified neurons was not simply due to cell loss due to death. There was no significant difference in the total neuronal density, although there was a significant trend toward neuronal loss at the 24 h time point when compared to the first time point before NMDA (p=0.02). (number of fields: 0h, 1h and 24h= 50; 30min, 3h and 12h= 70). Data are mean ± SEM, from two representative culture preparation; data collected from 50-70 fields of view from multiple coverslips. (**E**) Time-lapse sequence of images collected at the indicated times (hours: minutes) from a dendritic region of an actinified neuron expressing mApple-Ftractin to monitor live the F-actin reorganization during recovery following NMDA stress removal (at t=0) via MK-801 addition as above in “D”. Note not only the clearing of the accumulated F-actin within the dendrite shaft, but also the re-emergence of F-actin accumulation in spine-like protrusions. Scale bar, 8 µm.

### Spontaneous reversal of actinification upon stress removal

Despite the apparently non-dynamic nature of the F-actin induced by NMDA, the actinification filaments were able to spontaneously disassemble upon removal of NMDA. When cultures were incubated with NMDA for 5 min, followed by addition of MK-801 to prevent further NMDA receptor activation, the fraction of neurons in fixed cultures showing actinification dropped from about 60% to less than 20% within 1 hour (Fig 3D). By 3 hours only 8% of neurons remained actinified, and by 12 hours only 4% of neurons remained actinified. Over this same time frame there was no significant decrease in the number of live neurons, although there was a trend toward decreased neuronal viability by 24 hours (Fig. 3D). On average, the time for half-maximal recovery after a 5 min incubation with NMDA was approximately 30 min.

Time lapse imaging of individual neurons similarly documented that actinification was reversible, and that F-actin within the somatodendritic compartment spontaneously returned to a control-like distribution following arrest of the ongoing NMDA stress. Fig 3E shows a neuron expressing mApple-F-tractin, illustrating that the NMDA-induced accumulation of F-actin began to detectably clear from the dendrite shaft within 17 minutes of the addition of NMDA antagonists. Simultaneously, F-actin re-emerged in dendritic spines, thus returning to a qualitatively normal distribution within 1 hour (Fig 3E). These observations indicate that actinification persists only while the NMDA receptor hyperactivation persists, and that endogenous mechanisms support the spontaneous depolymerization of F-actin in the dendrite shaft/soma and the re-polymerization of F-actin in dendritic spines.

### Actinification is triggered by conditions that elicit neuronal swelling

Most of the early neuronal cell death induced during ischemia occurs via pathological cell swelling (also called cytotoxic edema) ^30, 31^. We observed that somatodendritic actinification occurred in parallel with swelling of the cell body and adjacent dendrite (Fig 4A). We therefore asked whether actinification was provoked by the same conditions that cause osmotic cell swelling, namely influx of both sodium and chloride, followed by water. First, we observed that replacement of extracellular sodium chloride by N-methyl-D-glucamine chloride completely prevented NMDA-induced actinification (Fig 4D), consistent with the hypothesis that sodium flux across the plasma membrane is critical for actinification. Recent studies demonstrated that voltage-gated chloride influx through the solute carrier family 26 member 11 (SLC26A11) ion exchanger drives neuronal Cl^-^ entry during glutamate-induced neuronal cell swelling ^10^. We found that actinification exhibited a similar pharmacological profile for chloride influx inhibitors as that reported for cytotoxic edema in brain tissue^10^. DIDS and GlyH-101, which inhibit Cl^-^ entry via SLC26A11 ^10^, significantly inhibited actinification, but neither bumetanide, which blocks the brain NKCC1 cation-chloride cotransporter, nor 5-nitro-2-(3-phenylpropylamino) benzoic acid (NPPB), which blocks volume regulated Cl^-^ channels and Ca^2+^-activated Cl^-^ channels, were effective in preventing actinification (Fig 4B). This indicates that actinification might require chloride entry via the same routes as reported for neuronal cell swelling.

**Figure 4.**
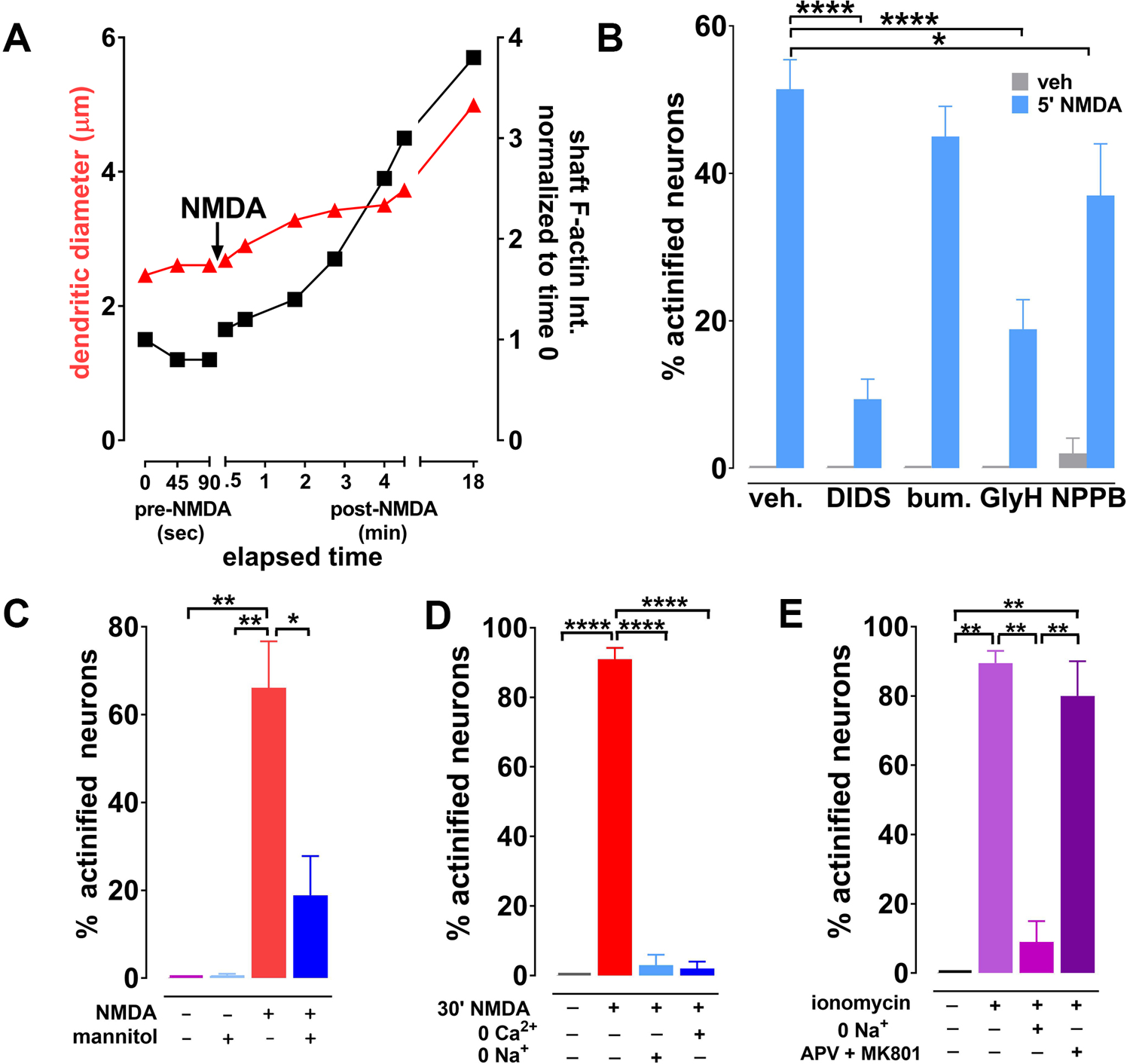
Evidence that actinification is triggered during cell swelling (cytotoxic edema). (**A**) Simultaneous quantification of dendrite diameter (*red triangles*) and F-actin intensity (*black squares*) within the dendritic shaft of a neuron imaged following addition of 50 µM NMDA at the indicated time. (**B**) Quantification of the effect of various chloride flux inhibitors on NMDA-induced actinification. Data are mean ± SEM from a minimum of two independent culture preparations; **** p<0.0001; *p<0.05; two-way ANOVA followed by Dunnett’s multiple comparison *post-hoc* test; number of coverslips per treatment group: veh= 6, DIDS=6, bumetanide (bum) = 6, GlyH= 3, NPPB=2 (drug abbreviations and molecular targets for each are described in Results and Methods. (**C**) Increasing extracellular osmolarity via addition of the cell impermeant sugar mannitol to the culture medium inhibits NMDA-induced actinification. Data are mean ± SEM from two independent culture preparations, with 4 coverslips in total per group; **p<0.01, *p<0.05; one-way ANOVA, followed by Tukey’s multiple comparison *post-hoc* test. (**D**) Removal of either sodium or calcium from the incubation medium prevents NMDA-induced actinification. Data are mean ± SEM from two independent culture preparations, with 4 coverslips in total per group; ****p<0.0001; one-way ANOVA, followed by Tukey’s multiple comparison *post-hoc* test. (**E**) Calcium-induced actinification requires extracellular sodium, but is not inhibited by NMDA receptor antagonists. Intracellular calcium was elevated using the calcium ionophore ionomycin (5 µM, 30 min) in the absence (-) or presence (+) of either extracellular sodium depletion or a cocktail of the NMDA antagonists APV and MK-801. Data are mean ± SEM from two independent culture preparations, with 4 coverslips in total per group; ** p<0.01; one-way ANOVA, followed by Tukey’s multiple comparison *post-hoc* test.

We next tested whether water entry was necessary for induction of actinification. Incubation of cultures with the non-cell permeable sugar mannitol, to reduce the osmotic driving force for water entry, indicated that swelling was indeed contributing to NMDA-induced actinification (Fig 4C).

### Calcium entry is necessary but not sufficient for actinification

NMDA receptors are highly permeable to calcium ions, and increased intracellular calcium is a trigger for many biochemical cascades elicited by NMDA receptor activation. Removal of extracellular calcium completely prevented neuronal actinification (Fig 4D). However, depletion of intracellular stores using thapsigargin (1 µM and 20 µM) had no effect in preventing actinification (*data not shown*). These data indicate that calcium influx across the plasma membrane is required to trigger actinification. However, calcium flux alone was insufficient, because although 10 min incubation with the calcium ionophore ionomycin induced substantial actinification, even in the presence of NMDA receptor antagonists, this effect was completely blocked when extracellular sodium was removed (Fig 4E). Collectively, our results indicate that sodium, chloride, and calcium influx are all required to induce actinification, and that water entry and consequent neuronal swelling is also a critical factor.

### Inverted formin 2 is required for actinification

We next turned to identifying the key actin mechanisms involved in catalyzing neuronal actinification. The polymerization of most actin filaments in cells are initiated via two distinct mechanisms. The Arp2/3 complex nucleates and elongates daughter filaments from the side of an existing actin filament, thereby forming branched F-actin networks, as seen in lamellipodia and dendritic spines^32–34^. Conversely, formin-driven F-actin polymerization induces formation of unbranched actin filaments, as seen in filopodia and other structures where straight filaments predominate^32, 35, 36^. We applied small molecule inhibitors of formin-mediated and Arp2/3-mediated actin polymerization, respectively. Only the formin inhibitor prevented NMDA-induced actinification (Fig 5A). This result is consistent with the ultrastructural observation that NMDA induced long, unbranched actin filaments (Fig 2D).

**Figure 5.**
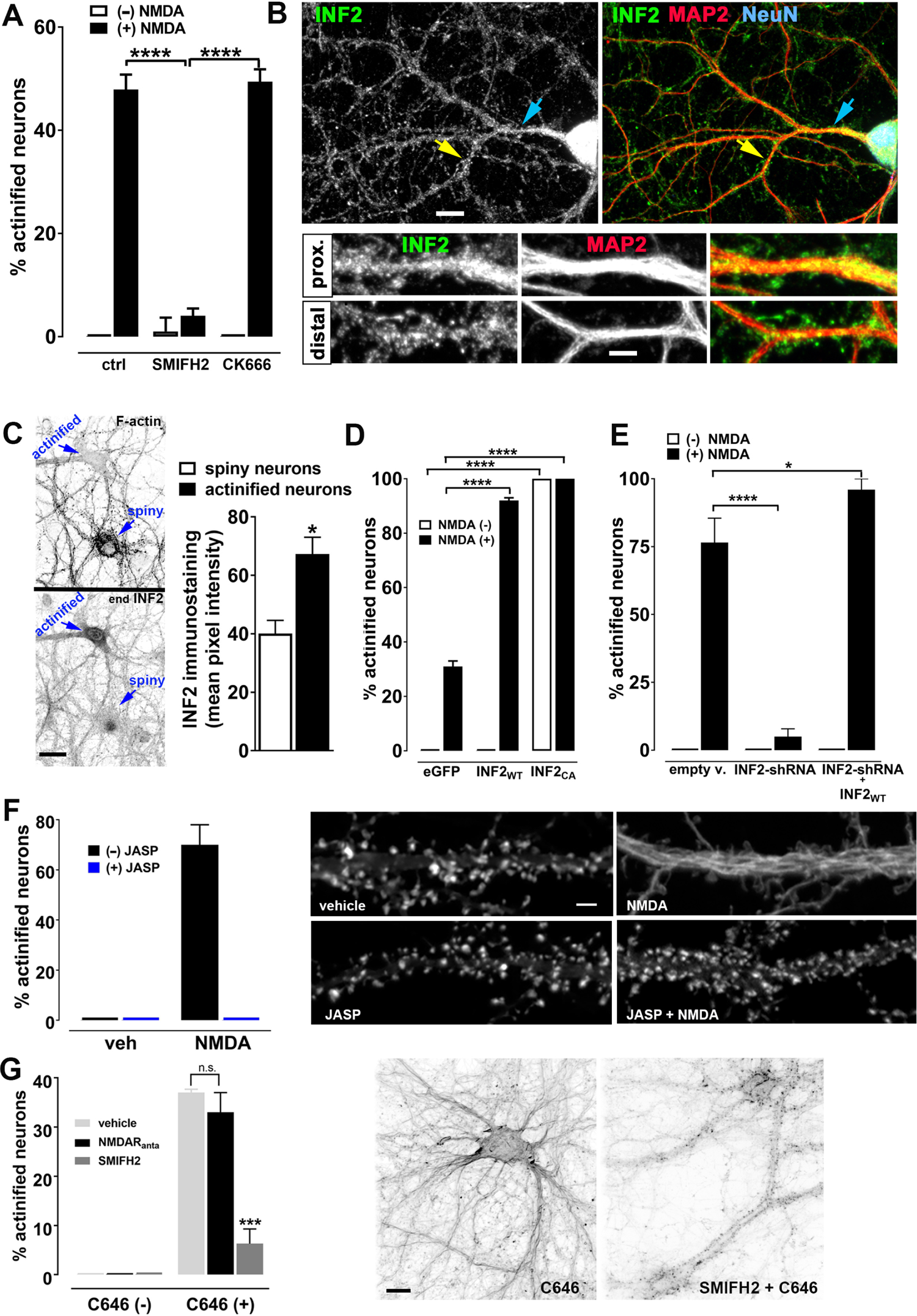
NMDA-induced actinification is driven by the formin INF2, and requires a decrease in acetylation and an initial F-actin depolymerization. (**A**) Inhibition of NMDA-induced actinification by the specific broad-spectrum formin inhibitor SMIFH2, but not the specific Arp2/3 inhibitor CK666. Data are mean ± SEM from two independent culture preparations, with 4 coverslips in total per group; ****p<0.0001; one-way ANOVA, followed by Tukey’s multiple comparison *post-hoc* test. (**B**) Representative images illustrating the distribution of endogenous INF2 (*green*) detected using immunostaining. Cultures were co-stained for NeuN (*blue*) and MAP2 (*red*), to identify neurons and their dendritic arbor, respectively. Arrows indicate dendritic regions that lie distal (*yellow*) and proximal (blue) to the cell soma, and are shown in higher magnification in the lower panels. Note the proximal-to-distal gradient of INF2 immunostaining intensity, with a higher overall signal within the soma and proximal dendrite. Scale bar, 12 µm (*upper* panels), 5 µm (zoomed *lower* panels). (**C**) Neurons that actinify quickly in response to NMDA exhibit higher concentrations of endogenous INF2. Selected images from a culture incubated for 5 min with 50µM NMDA displaying actinified and non-actinified (“spiny’) neurons within the same field-of-view; the earliest responders typically express higher endogenous levels of INF2 immunoreactivity, as quantified on the *right*. *Blue arrows* point to the neuronal cell bodies, where INF2 staining intensity was quantified across multiple samples. Scale bar: 20 µm. Data are mean ± SEM from two independent culture preparations, with 4 coverslips in total per group; *p<0.05; Unpaired Student’s t-test, two tailed. (**D**) Ectopic expression of INF2 sensitizes neurons to NMDA-induced actinification. Note that the presence of wildtype INF2 (INF2WT) did not induce actinification in the absence of NMDA, but significantly increased the fraction of neurons that actinified in the presence of 5 min 50µM NMDA, compared to the control group transfected with eGFP. In contrast, expression of a constitutively active form of INF2 (INF2CA) induced actinification in both the absence and presence of NMDA in all neurons. Data are mean ± SEM from 4 independent culture preparations, with a minimum of 2 coverslips in total per group; ****p<0.0001; two-way ANOVA, followed by Dunnett’s multiple comparison *post-hoc* test. (**E**) Silencing of INF2 gene expression via shRNA prevents NMDA-induced actinification. The following groups of transfected cultures were incubated in the absence (-) vs. presence (+) of NMDA: Control (empty vector); INF2 knockdown by shRNA; rescue by ectopic INF2 (INF2shRNA + INF2-wt resistant to knockdown; both CAAX and non-CAAX variants of INF2-wt were tested, and as we observed no statistical difference in the degree of rescue between them we combined the data into a single group). Data are mean ± SEM from two independent culture preparations, with 2-5 coverslips in total per group; *p<0.05, ****p<0.0001; two-way ANOVA, followed by Dunnett’s multiple comparison *post-hoc* test within NMDA(-) and NMDA(+) samples. (**F**) NMDA-induced actinification is completely prevented by F-actin stabilization. GFP-Ftractin expressing neurons were incubated in the absence or presence of NMDA (50 µM, 5 min) in the absence or presence of jasplakinolide added simultaneously (JASP, 4 µM) Data are represented as mean ± SEM; n=2. (**I**) Selected proximal regions of the main dendrite of neurons treated as indicated in the image panels. Scale bar, 4 µm. (**G**) Evidence that protein acetylation regulates actinification, despite NMDA receptor blockage by a cocktail of NMDAR antagonists (APV + MK801); acetylation was increased globally by incubation for 30 min with the broad spectrum acetyltransferase inhibitor C646. As shown by the graph and corresponding images, C646-induced actinification, similar to NMDA-induced actinification, requires formin activity, since both are blocked by the formin inhibitor SMIFH2. Data are mean ± SEM from 3 independent culture preparations, with 6 coverslips in total per group; ****p<0.0001, n.s.= not significant; two-way ANOVA, followed by Tukey’s multiple comparison *post-hoc* test. Scale bar, 12 µm.

Formins constitute a large superfamily of molecules, with fifteen mammalian formin genes identified to date^37^. Our attention was drawn in particular to inverted formin 2 (INF2), a member of the diaphanous subclass of formins. We found that immunoreactivity for INF2 is present throughout the somatodendritic domain of cultured hippocampal neurons, and distributes in a punctate fashion within both proximal and distal dendrites (Fig 5B). No such staining was observed when endogenous INF2 was depleted from individual neurons via RNA interference (Suppl Fig. S6). We observed a gradient of INF2 immunoreactivity within the dendritic arbor, with highest intensity in the soma and proximal dendrites, which diminished gradually toward the more distal dendrites (Fig. 5B). This distribution gradient in control neurons resembles that of actinification itself.

Interestingly, we observed some variability in the relative concentrations of INF2 immunoreactivity across neurons within the same culture. We treated cultures for 5 min with NMDA (a time when typically 35-60% the neurons have become actinified) and quantified the level of immunoreactive staining for INF2 in the soma of actinified neurons versus non-actinified neurons (Fig 5C). We observed a significantly higher degree of INF2 staining in the neurons that were the “early responders” – i.e., those that underwent actinification at this early time point. This bias in endogenous INF2 level implies that the concentration of INF2 may at least partly determine the strength or timing of the actinification response in individual neurons.

To test this hypothesis directly, we transfected neurons with a constitutively active form of INF2 and observed that neuronal soma and dendrites showed actinification even in the absence of NMDA exposure (Fig. 5D). More importantly, neurons transfected to ectopically express a wildtype form of INF2 showed no increase in actinification in the absence of NMDA, but showed a dramatically enhanced actinification response when stimulated with NMDA for 5 min, with nearly all the neurons becoming actinified within this short time frame (Fig 5D).

Conversely, silencing of endogenous INF2 expression using an shRNA completely blocked the actinification of the neuron, an effect that was rescued in the presence of an shRNA-resistant form of wildtype INF2 (Fig 5E). Both CAAX and non-CAAX variants of INF2-wt ^38^ were tested, and as we observed no statistical difference in the degree of rescue between them we combined the data into a single group. Together, these observations implicate INF2 as a critical driver of NMDA-induced actinification. They also indicate that INF2 activity normally remains low under control conditions regardless of INF2 concentration, and must be induced by a stimulus, since overexpression of wildtype INF2 had no detectable effect on actinification under resting conditions.

### A role for elevated G-actin, but perhaps not for deacetylases in NMDA-induced actinfication

Recent studies showed that INF2 is maintained in an inactive conformation in cells^39^ and is then activated by stimuli that raise intracellular calcium ^40–42^. One mechanism for this activation might be an increase in G-actin levels, since INF2 is known to be stimulated by elevated actin monomer concentrations ^39^. We therefore hypothesized that the G-actin released during NMDA-induced depolymerization of spine F-actin might activate INF2. In support of this hypothesis, adding jasplakinolide during the 5 min NMDA incubation completely prevented both spine F-actin disassembly and dendritic actinification (Fig 5F).

INF2 has also been proposed to be negatively regulated by its binding to a complex consisting of cyclase-associated protein (CAP) and acetylated-G-actin. This complex is disrupted when G-actin becomes deacetylated, thereby allowing activation of INF2 ^43, 44^. We observed that incubation of cultured neurons with the acetyltransferase inhibitor C646 strongly induced actinification, an effect that was not blocked by NMDA antagonists, consistent with a role for acetylation in regulating actinification (Fig 5G). Notably, the C646-induced actinification was almost completely blocked by preincubation with the formin inhibitor SMIFH2 (Fig. 5G), similar to actinification induced by NMDA (Fig 5A), suggesting they are mediated by the same INF2-dependent mechanism. We therefore hypothesized that INF2 activity might become induced in response to NMDA via the activation of cytosolic deacetylases (which are typically called histone deacetylases, or HDACs, even though it is now known that they can deacetylate numerous cytosolic substrates). As shown in Table 1, pre-incubation with several inhibitors of Class 1 or Class 2 HDACs modestly reduced NMDA-induced actinification, but none were robustly effective, including the HDAC6 inhibitor tubastatin, which blocked INF2 activity in non-neuronal cells ^44^. Moreover, combinations of multiple inhibitors qualitatively showed little or no additive effect toward inhibiting actinification. Therefore, while our data suggest that de-acetylation is a factor in regulating neuronal actinification, the precise mechanisms that lead to INF2 activation in response to NMDA-induced cellular edema might not require deacetylase activity.

**Table. 1.** HDAC inhibitors (HDI) and their efficacy in reducing NMDA-induced actinification.

| HDAC inhibited | Treatment<br>(1h HDI + 5min NMDA) | % Reduction<br>in actinification (n) |
| --- | --- | --- |
| **Class I + II** | SAHA (1 μM) | 40 ± 6 (5) |
| **Class I** | Santacruzamate (0.3 μM) | 33 ± 3 (3) |
| **Class I** | CI994 (1 μM) | 30 ± 4 (3) |
| **Class I** | FK228 (1 μM) | 36 ± 3 (6) |
| **Class II** | Tubastatin (5 μM) | 21 ± 6 (6) |
| **Class II** | Tubacin (10 μM) | n.d.q. |
| **Class III** | Nicotinamide (2 mM) | n.d.q. |
| **Class III** | Ex527 (2.5, 5, 30, 100 μM) | n.d.q. |
| **Class III** | anti-SRT5 peptides (20 μM) | n.d.q. |
| **Class III** | Suramin (100 μM) | n.d.q. |

Cultured hippocampal neurons were incubated for 1h with HDAC inhibitors (HDI) before incubating the cells for 5 min with 50 μM NMDA. Concentrations below and above the ones listed were also tested to reach either no or toxic effect. *n.d.q.* = *no inhibition detected in qualitative analyses*; n = number of independently repeated experiments.

### NMDA-induced accumulations of actin in the shaft are distinct from cofilin-actin rods

Previous studies have reported that excess neuronal glutamate or hypoxic stress induce the accumulation of aberrant F-actin containing structures called cofilin-actin rods, which are defined by the concentrated presence of the actin severing protein ADF/cofilin along bundles of F-actin^45^. Such actin rods are not detectable by phalloidin staining, since the prominent binding of cofilin prevents the binding of phalloidin in such filaments ^46^. Four lines of evidence indicate that the actinification of the dendrite compartment is a different process than the formation of cofilin-actin rods. First, unlike cofilin-actin rods, the actin filaments we observe are clearly labeled by phalloidin. Second, these filaments do not contain high concentrations of cofilin immunoreactivity (Suppl Fig S7A). Third, the spatiotemporal dynamics of actinification and the emergence of cofilin-actin rods differ qualitatively. Cofilin-actin rods reportedly appear in dendrites only after 30 min or more of continuous exposure to glutamate or NMDA^45^. Although we can detect cofilin-actin rods after a lengthy exposure of our cultures to glutamate (Suppl. Fig. S7B1 and B2), typically they appear mainly in the distal dendrites and along axons surrounding the dendritic arbor. In contrast, actinification occurs within less than 5 minutes and is preferentially detected in the cell body and the proximal dendrites. Finally, even cofilin rods induced without NMDA by ectopic expression of wild type cofilin together with one of its activating phosphatases, chronophin do not promote neuronal actinification (Suppl Fig S7C). Together, these observations strongly argue that somatodendritic actinification is a novel process that is distinct from cofilin-actin rod formation.

### Actinification is a pro-survival response

We next investigated the functional impact of INF2-dependent actinification. Because the distribution of actin filaments spontaneously returned to control values after cessation of the stressful stimulus, we reasoned that actinification might be a pro-survival response. To test this hypothesis, we incubated cultures with NMDA for 1 or 4 hours and compared neuronal survival using the VivaFix cell viability assay in neurons that were either transfected with shRNA against INF2 or with an empty vector (Fig 6B). For both conditions, neurons were co-transfected with eGFP to identify transfected cells, and we quantified the percentage of transfected neurons that took up the VivaFix dye as an indicator of cell death (Fig 6A). We observed that after either a 1 hour or 4 hours incubation with NMDA the prior silencing of INF2 approximately doubled the fraction of non-viable neurons.

**Figure 6.**
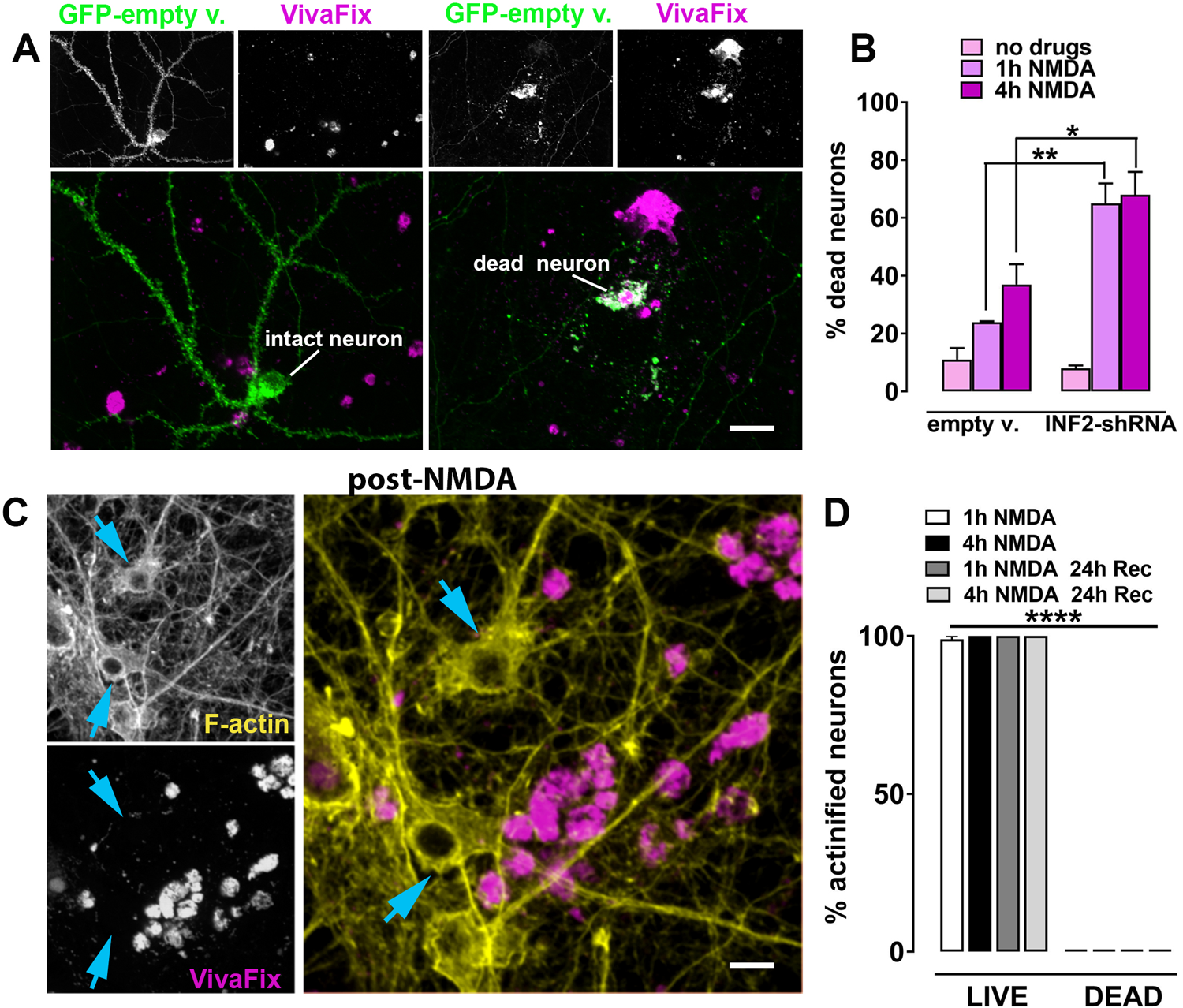
INF2 silencing enhances NMDA-induced cell death in vitro. Hippocampal neurons expressing either a control empty vector or its corresponding INF2-shRNA were incubated in the absence or presence of NMDA for 1 h or 4 h, as indicated, prior to fixation and co-staining with Alexa fluor 568-phalloidin and the VivaFix 649/660 reagent. (**A**) Representative images of live, intact neurons (those that exclude the red VivaFix dye) vs. dead neurons (those that incorporate the VivaFix dye). GFP-empty vector (*green*) was used in live imaging mode to evaluate neuronal morphology (e.g., intact vs. fragmented). Cell viability was assessed using the VivaFix assay (*magenta*). Scale bars, 28 µm. (**B**) Silencing of INF2 is associated with a significant enhancement of time-dependent NMDA-induced neuronal cell death. Data are mean ± SEM from 3 independent culture preparations; *p<0.05; **p<0.01; two-way ANOVA, followed by Tukey’s multiple comparison *post-hoc* test. (**C**) Representative images of neurons incubated for 1 h with NMDA and fixed immediately, illustrating that NMDA-induced actinified neurons (*yellow*=phalloidin) do not take up the cell death marker VivaFix (*magenta*). Scale bars, 14 µm. (**D**) Quantification of the live vs. dead fraction of actinified neurons among the neurons transfected with the control, empty vector revealed a binary fate for neurons incubated for 1-4 hours with NMDA: those neurons that exhibited actinification never took up the VivaFix dye; those that did take up the VivaFix dye were never observed to be actinified, whether fixed and stained immediately after the indicated NMDA incubation, or fixed after an additional 24 hours when the 1 or 4 hour NMDA incubation was followed by a potential “recovery period” imposed by the addition of a cocktail of APV and MK-801 to prevent ongoing excitotoxic stress. Data are mean ± SEM from two independent culture preparations; **** p<0.0001; two-way ANOVA, followed by Tukey’s multiple comparison *post-hoc* test.

By 24 hours after incubation with NMDA for 1-4 hours, the vast majority of neurons had died. However, a very small number of live neurons were still reliably detected (i.e., they excluded the VivaFix dye), even after this strong excitotoxic stimulus. Remarkably, 100% of these late-surviving neurons displayed actinification (Fig 6C-D). Conversely, none of the neurons that were identified as dead showed actinification (Fig 6C-D), and, indeed, phalloidin staining was depleted in dead neurons.

### Formin inhibition exacerbates cell death after stroke *in vivo*

We next asked whether INF2-driven actinification plays a role in neuronal survival following ischemic stroke. Focal stroke was induced using single vessel photothrombosis in mouse cortex. Brains were fixed 4 hours after induction of vessel occlusion, and Fluorojade C was used as a marker of early cell death in post-fixed histological sections to evaluate the effect of inhibiting formin activation using the compound SMIFH2. SMIFH2 or vehicle were applied 4 hours prior to stroke induction using an agar-saturated plug gently placed over the thinned skull. We found that pre-treatment with the formin inhibitor induced a near doubling of cell death after stroke, compared to that observed with vehicle treatment (Fig 7A-C; Suppl Fig S8). Neither vehicle nor SMIFH2 induced detectable FluoroJade C staining in the absence of stroke.

**Figure 7.**
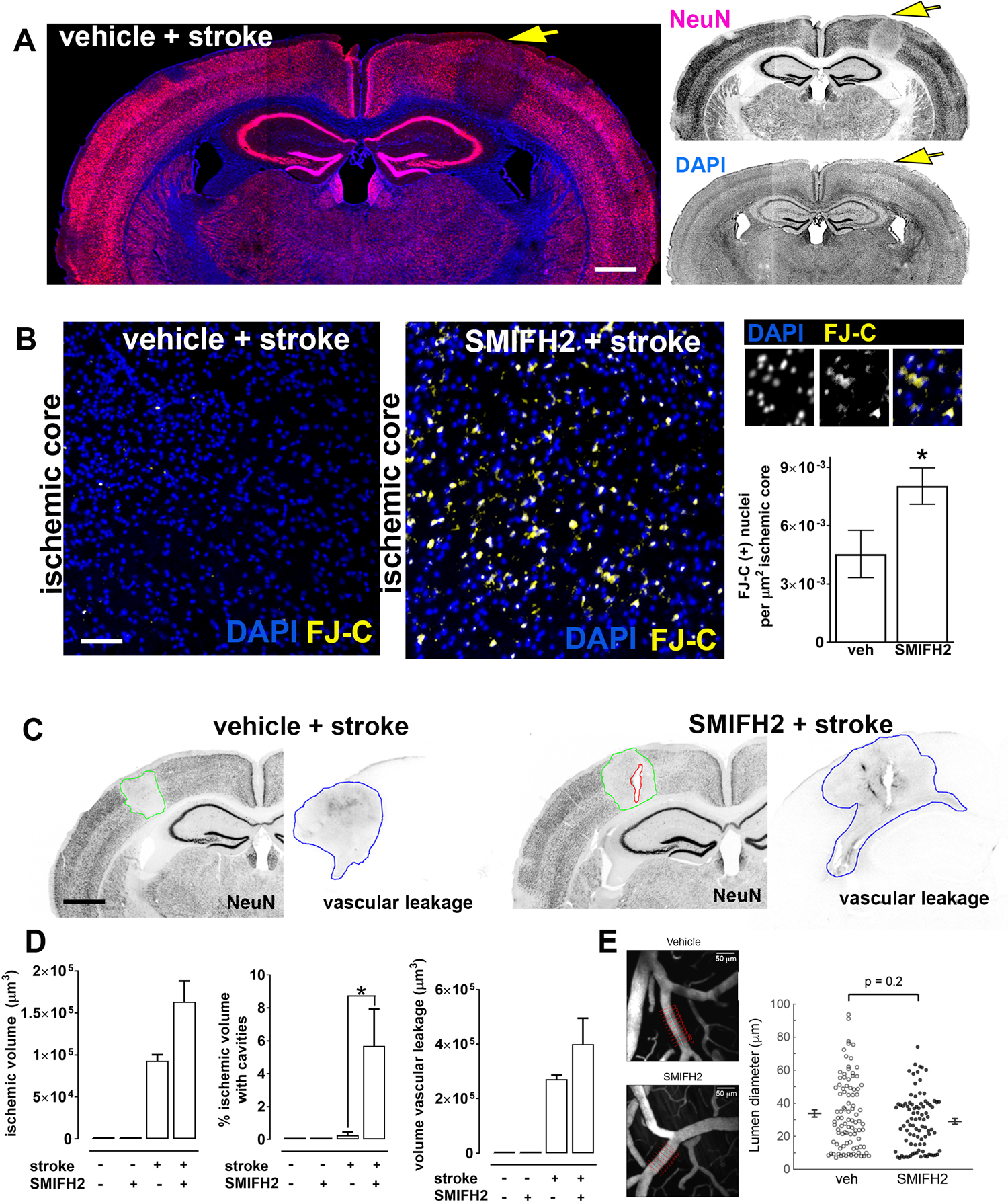
Formin inhibition enhances stroke-induced cell death *in vivo*. (**A**) Coronal mouse brain section illustrating the loss of NeuN immunoreactivity within the small somatosensory region receiving a single vessel photothrombotic stroke (region indicated by yellow arrow). Prior to the stroke the DMSO vehicle (the control for the SMIFH2 drug) was delivered to the infarct zone for 30 min by passive diffusion, as described (see Methods). Scale bar, 850 µm. (**B**) Inhibition of formin activity by SMIFH2 increases early cell death induced by stroke. Representative cortical regions within the ischemic core are shown. Nuclei are labeled with DAPI in *blue*, and the cell death marker FluoroJade-C (FJ-C) in *yellow*. Panels at *right* show enlarged regions illustrating FJ-C nuclear labeling colocalized with DAPI. Quantification of the density of FluoroJade-C-positive (+) nuclei within the ischemic core region defined by the loss of NeuN signal. Data are mean ± SEM; * p<0.05; Student’s t-test, unpaired, two-tailed; number of animals: veh + stroke = 7, SMIFH2 + stroke= 8. Scale bar: 70 µm. (**C**) Representative images of brain sections stained for the neuronal marker NeuN and for mouse anti-IgG to assess the extent of vascular leakage following stroke. Stroke was induced 30 min after vehicle (DMSO) or the formin inhibitor (SMIFH2) were passively diffused into the ipsilateral hemisphere. Scale bars, 800 µm. (**D**) Characterization of ischemic lesion in the presence of vehicle (DMSO) vs. SMIFH2; mice receiving the same drug delivery but no infarct were used as controls for the drug delivery manipulations. Brain section were blinded to treatment group, and stained to evaluate infarct volume (based on NeuN signal loss), the fraction of the ischemic volume subsumed by cavitations (i.e., holes in the brain parenchema), and vascular leakage (based on immunoreactivity for mouse IgG). Data are mean ± SEM. *p<0.05; one-way ANOVA, followed by Dunnett’s multiple comparisons *post-hoc* test vs. veh+stroke); number of brains: veh./no stroke = 3, SMIFH2/no stroke = 3, veh + stroke = 7, SMIFH2 + stroke = 8. (**E**) *Left,* Representative *in vivo* two-photon imaging of cortical pial vessels from vehicle-treated (*upper*) and SMIFH2-treated (*lower*) mice. Red lines placed across the width of a pial arteriole denote the position of diameter measurements that were averaged to provide a single data point. Scale bars, 50 µm. *Right*, Scatter plot of pial arteriole diameters from vehicle and SMIFH2-treated groups, including 93 arterioles over 6 vehicle-treated mice, and 90 arterioles over 7 SMIFH2-treated mice; mean and SEM are indicated adjacent to each scatter plot; no significant differences were detected between groups, statistical analysis performed by Wilcoxon test.

Quantification of the ischemic volume indicated that formin inhibition induced a non-significant trend toward increased infarct volume at 4 hours post-occlusion (Fig 7C and D). Moreover, we observed a substantial increase in apparent damage to cortical tissue within the core of the infarct, with variably sized cavities appearing after strokes induced in the presence of SMIFH2. Such cavitation was never observed in non-infarcted brain regions nor in vehicle treated brains either with or without stroke. Despite these indicators of enhanced tissue damage following stroke, SMIFH2 did not induce a significant increase in the leakage of immunoglobulin from compromised blood-brain barrier following vessel occlusion (Fig 7C and D). In addition, direct measurement of unoccluded arteriole diameters in the vicinity of the occluded vessel showed no differences between vehicle or SMIFH2 treatment (Fig 7E), indicating that the drug did not alter local blood flow dynamics prior to stroke induction. These data indicate that INF2 normally helps protect ischemic brain tissue during the acute phase of infarct development.

## DISCUSSION

Cellular pro-survival responses engage multiple subcellular events to enable cells to endure periods of transient stress ^47^. Excitable cells like neurons and cardiomyocytes are especially vulnerable to osmotic stress because perturbed flux through various ion channels can lead to osmotic imbalance. Swelling in these cells and tissues endangers the survival of the organism. Due to the critical role of ATP-dependent ion pumps in osmoregulation, catastrophic reduction in cellular ATP throws such homeostatic mechanisms into disarray. Here we show that the neuronal actin cytoskeleton undergoes a rapid and dramatic reorganization in response to ischemia or excitotoxic levels of glutamate -- conditions that trigger cytotoxic edema. This neuronal response is reversible and pro-survival.

Our data suggest a model in which a strong influx of sodium and chloride, and subsequent water entry (i.e., the key drivers of cell swelling), along with an influx of calcium ion, are necessary and sufficient to induce a fundamental reorganization of the actin cytoskeleton within the somatodendritic compartment of neurons. This convergence of ion influx leads to the activation of the diaphanous family formin INF2. We postulate that INF2 is required either to nucleate new filaments or to elongate existing short filaments within the soma and dendrite. The identification of a formin-based mechanism for actinification is consistent with several lines of evidence, including the long, unbranched filaments observed using both light and electron microscopy, the complete inhibition of actinification by the formin inhibitor SMIFH2, and the complete prevention of actinification by genetic silencing of INF2, which was rescued by ectopic expression of INF2. Interestingly, although we could observe actinification in live cells using multiple fluorescent reporters of actin filaments (Lifeact-mRFP, mApple-Ftractin, and the small molecule probe SiR-actin), we were unsuccessful in observing actinification using GFP-actin. This also is consistent with a formin based mechanism, since various reports indicate that large moiety-terminal tags on G-actin interfere with formin-dependent F-actin assembly ^48–52^.

Importantly, several lines of evidence establish that the actinification phenomenon is distinct from the formation of cofilin-actin rods, which can also form in response to excess glutamate ^45^. First, the spatial and temporal features of these events are different, with actinification occurring rapidly within 5 minutes and predominantly in the soma and proximal dendrites, while cofilin-actin rods reportedly appear after at least one hour, preferentially in distal dendrites ^53^, an observation that we confirmed in our own culture system. Secondly, we found that cofilin immunoreactivity did not colocalize with the actinified filaments as would be expected for cofilin-actin rods. Finally, multiple means of preventing or inducing cofilin activation neither prevented or favored actinification. The accumulation of F-actin within the dendrites also did not resemble typical actin stress fibers, since it was not prevented by blebbistatin or by the Rho-kinase inhibitor fasudil (data not shown). We therefore conclude that the actin filaments that accumulate within the somatodendritic compartment following osmotic stress in neurons are distinct, and characterized here for the first time.

The precise mechanisms by which swelling and calcium entry converge to activate INF2 require further investigation. Higgs and colleagues have shown that, in non-neuronal cells, an inactive conformation of INF2 is maintained via binding of a complex of lysine acetylated G-actin bound to cyclase-associated protein (CAP) to the INF2 DID and DAD domains, respectively, and that INF2 becomes activated when HDAC6 activity deacetylates G-actin ^44^. In our studies of primary neurons, however, none of the various deacetylase inhibitors we tested, including the HDAC6 inhibitor tubastatin, robustly inhibited NMDA-induced actinification. Nevertheless, some mechanism involving acetylation/deacetylation activity does seem to influence actinification driven by INF2, since the compound C646, which broadly inhibits acetyltransferases, by itself caused actinification, even in the presence of NMDA antagonists. This effect of C646 was blocked in the presence of the formin inhibitor SMIFH2, indicating it may act via INF2, similar to NMDA. Taken together, we cannot rule out that NMDA induces INF2 activity via a pathway involving deacetylation, but our results suggest that NMDA possibly activates INF2 through alternative mechanisms.

Studies have demonstrated *in vitro* and in cells that elevated actin monomer concentration can compete with INF2 autoinhibition in addition to its role as nucleation substrate^39, 54^. We postulate that the abrupt rise in G-actin driven by NMDA-induced depolymerization of spine actin generates a burst of soluble monomers that may facilitate the activation of INF2. Indeed, prevention of F-actin disassembly by jasplakinolide completely blocked NMDA-induced actinification (Fig 5F).

The filaments formed during actinification appear to exhibit an unusually slow turnover. The failure of GFP-tagged forms of exogenous actin to participate in actinification precluded our ability to directly quantify actin turnover rates following NMDA. However, when latrunculin A was applied to sequester G-actin after actinification had occurred, we observed no enhanced clearance of actinified filaments for up to 2 hours, suggesting that there is very little turnover of these filaments in the continued presence of NMDA. We therefore conclude that the half-time for filament turnover probably exceeds ∼1 hour, meaning they are highly stable. Despite this remarkable apparent stability, the filaments are able to spontaneously disassemble upon cessation of NMDA receptor activation. Although detailed characterization is required, we estimate that the half-time for filament disassembly (when NMDA antagonists are applied after 5 min of NMDA) is on the order of 15-45 minutes. Interestingly, F-actin also reassembled in dendritic spines during the same time frame for F-actin disassembly in the dendrite shaft, a phenomeon that also deserves further investigation.

Initially, we had assumed that the massive reorganization of F-actin represents an early step in excitotoxic neuronal cell death. Indeed, previously we reported that NMDA-induced F-actin reorganization was attenuated by lithium ^14^, which has been implicated in neuroprotection ^55^. However, as discussed below, we determined that actinification is pro-survival. Given that actin filament assembly is typically an energy-consuming process, it is curious that neurons would engage in large-scale F-actin polymerization during a time of cell stress, especially during hypoxia when ATP is in short supply. The ATP-dependence of actinification remains to be determined; however, polymerization of ADP-actin into filaments has been described ^56, 57^.

The reversible nature of somatodendritic actinification suggested that it might be beneficial to the cell. Subsequent experiments convincingly demonstrated the pro-survival nature of this pathway (Fig 6B). Prolonged incubation with NMDA induced a portion of cells to die within 1-4 hours, probably via swelling and necrosis rather than apoptosis, due to its rapidity. We chose lengthy exposures to NMDA for these experiments in order to maximize cell death induction, since most neurons incubated with NMDA for shorter periods do not succumb to stress immediately, but rather over 24-48 hours ^58^. Note that even after 1 hour of continuous exposure to NMDA only ∼25% of control, empty vector-transfected neurons were dead at this time point, and only ∼35% were dead after 4 hours of continuous exposure to NMDA (Fig 6B). Silencing of INF2 significantly enhanced this rapid neuronal death.

Moreover, pharmacological inhibition of formin activity *in vivo* confirmed that blocking this pathway significantly worsened ischemic infarct severity. Given that at early times (i.e., hours) following a stroke most of the neuronal cell death that generates the infarct is mediated by pathological cell swelling ^31, 59^, we postulate that INF2-mediated actinification attenuates the effects of cell swelling and reduces cell death in the early stages after a stroke.

Consistent with this hypothesis, we observed the appearance of small cavities within the core of the infarct in our experimental model of stroke, but only when formin activity was inhibited (Fig 7). Cavitation within infarct zones has been described previously as cavities which appear many days or weeks after a stroke, evolving from a cystic, fluid filled core ^60, 61^. The early appearance of tissue cavities was therefore unexpected, and this observation is worthy of follow up investigation. One possibility consistent with our model is that blockage of formin activity renders neurons so susceptible to edema that they undergo cytolysis, leading to rapid tissue damage and cavitation in the acute ischemic core.

The pro-survival function of INF2-mediated actinification might be selective for neuronal edema or related types of osmotic stress. We observed that actinification required a convergence of calcium entry together with an ionic imbalance and water entry. This may imply that actinification selectively protects neurons from cytotoxic edema, but not other stressful conditions.

It will be of interest to determine whether actinification is a relevant pro-survival mechanism in widely-occurring injury conditions involving glutamate receptor overload, including traumatic brain injury and seizures in addition to hypoxia/ischemia.

Previous studies have also implicated INF2 in mechanosensitive responses of cells in culture, but the *in vivo* relevance of these responses has not been explored. A study in XTC cells demonstrated mechanosensitive activation and processive F-actin polymerization by diaphenous formins, including INF2 ^62^. A study in NIH 3T3 cells showed that INF2 mediated the formation of a perinuclear actin rim in response to mechanical stress or calcium ionophore, but did not determine the function of this actin rim structure ^41^. Another study similarly reported that INF2 mediates a transient response to cell damage or strong calcium entry, with actin polymerization occurring along the ER, simultaneous with actin depolymerization at the cell periphery, a response the authors termed “calcium-mediated actin reset” ^42^. Because the temporal dynamics, sodium and chloride dependence, and other key features of the actin polymerization events described in these prior studies appear to differ from the neuronal actinification we show here, further investigations are needed to examine the relationship among these various cytoskeletal responses. Nevertheless, it is worth postulating that INF2-driven actin polymerization may function in a general pro-survival capacity to protect many types of cells from a variety of mechanical stressors, including swelling during ischemic, hypoxic, or osmotic episodes.

## Acknowledgements

We are grateful to the laboratory of Mana Parast (UC San Diego) for use of their hypoxic chamber, and Christian A. Olsen for the generous gift of specific SRT5 inhibitors. We thank Vaidehi Gupta for generating dissociated neuronal cultures and other support, and Agnieszka Brzozowska-Prechtl for brain sectioning. We thank David Kleinfeld (UC San Diego) and Halpain lab members for helpful discussions. We greatly appreciate the following peerless undergraduate lab volunteers for conducting blinded data acquisition and analysis and general lab support: Kyra Rashid, Jeremy Aung, Liam Huber, Shyam Patel, Youjia Guo, Molly Thapar, Huanqiu Zhang. We thank the UCSD Nikon Imaging Center for access to the NIS Elements analysis software. This research was supported by grants from the U.S. National Institutes of Health, grant MH087823 (S.H.); and the Air Force Office of Scientific Research, grant FA9550-18-1-0051 (S.H.); NIH grant R35 GM140832 (T.M.S.); NIH grants NS106138 and NS097775 (A.S.).

## Author contributions

B.C. and S.H. designed the study and wrote the manuscript with input from all authors. S.J. and T.S. conducted the electron microscopy experiment. A.S. conducted the *in vivo* stroke experiments and analysis of arteriole diameters. B.C. conducted all other experiments. Y.Y. carried out initial experiments that identified neuronal actinification. U.M. helped with the Airyscan acquisition. H.H. provided INF2 constructs and advised on their use. M.L. generated the mApple-INF2-shRNA.

## Declaration of interests

The authors declare no competing interests.

## Methods

Detailed methods are provided in the online version of this paper and include the following:

### KEY RESOURCES TABLE

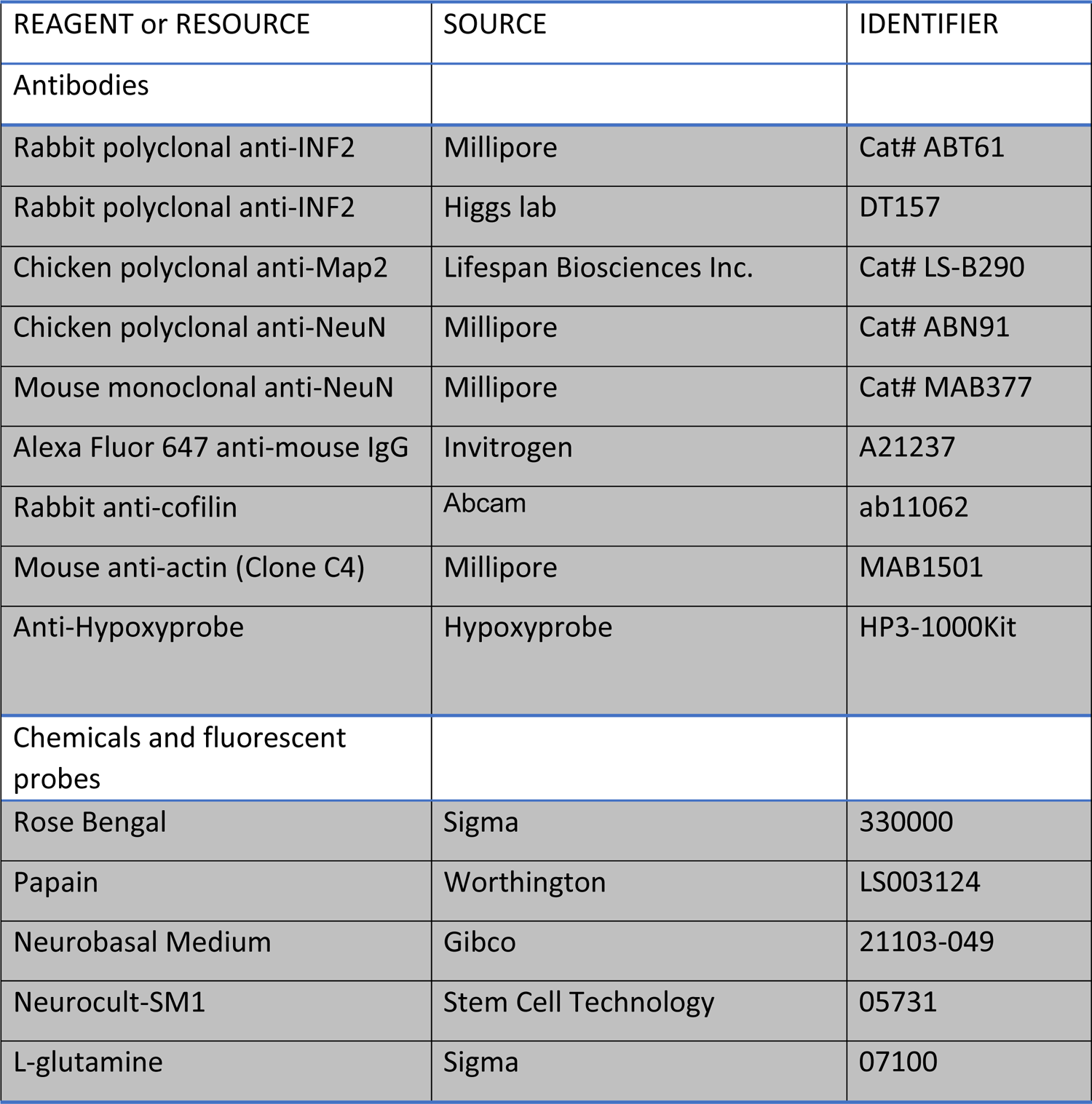
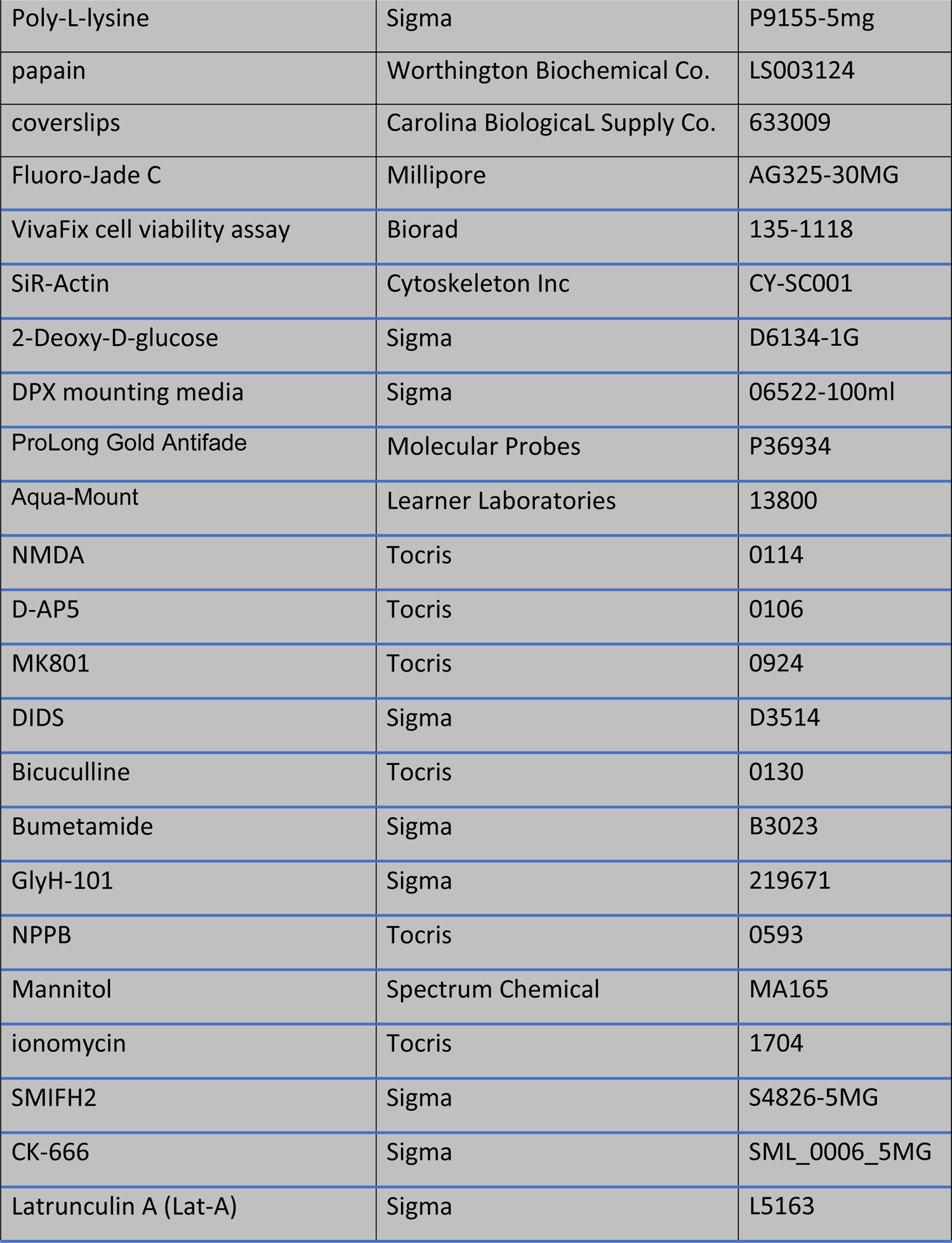
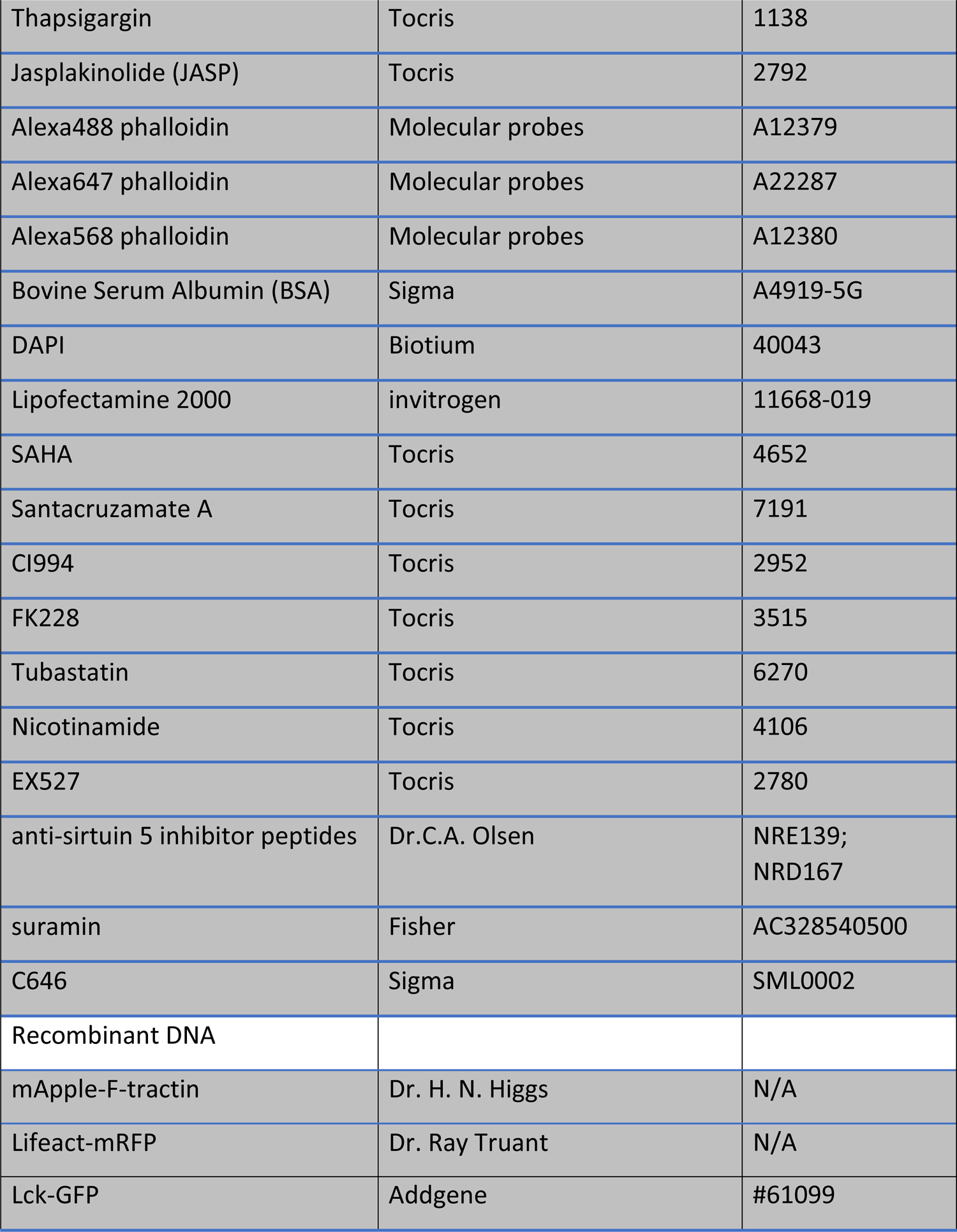
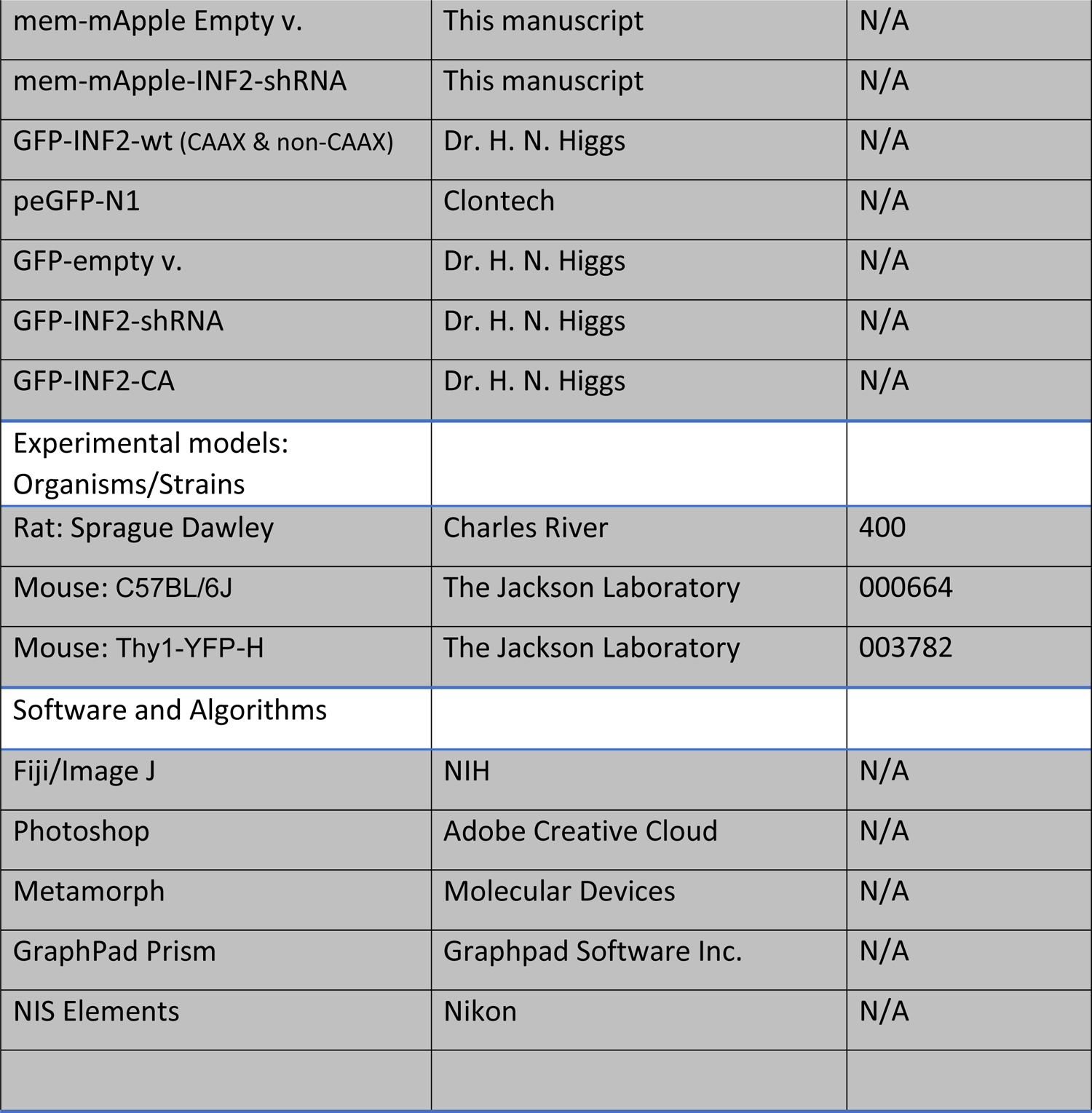

### CONTACT FOR REAGENT AND RESOURCE SHARING

Further information on procedures and requests for reagents should be directed to and will be fulfilled by Dr. Barbara Calabrese, Dr. Shelley Halpain, Dr. Andy Shih, Dr. Uri Manor, Dr. Tatyana Svitkina, or Dr. Henry N. Higgs.

### EXPERIMENTAL MODELS

#### Primary rat neuronal culture

Rat hippocampal neurons were isolated according to Calabrese and Halpain ^1^. In brief, hippocampi were dissected from brains of Sprague-Dawley rat (Charles River) female and male embryos at embryonic day 19 and dissociated into individual cells by incubating in a papain-containing solution. The cells were then washed and plated either on poly-L-lysine-coated (100 µg/ml) glass coverslips or on PEI (1:100 in borate buffer) + Laminin (20 µg/ml) 24 well ibiTreat (ibidi) at a density of 500 cells/mm^2^ in Neurobasal Medium (Gibco) supplemented with Neurocult SM1 neuronal supplement (Stem Cell Technology) and 0.5 mM L-glutamine (Sigma).

Neurons were usually transfected at 21 days *in vitro* (DIV) using Lipofectamine 2000. Cells were fixed or used for live cell-imaging experiments 1 to 2 days post-transfection, or 7-10 days post-transfection for the shRNA experiments.

#### In vivo model

One-to-two months old C57BL/6J and Thy1-YFP-H mice, both male and female, were received from Jackson laboratory and used for the *in vivo* experiments described in Fig 1D, E & F, Suppl. Fig S1-S5 and in Fig 7 and Suppl. Fig S8 respectively. The Thy1-YFP-H mice, as previously reported, have YFP expression restricted mainly to subsets of layer V neurons facilitating the identification of dendrites and dendritic spines ^2^.

## METHOD DETAILS

### Plasmid construction for INF2 RNA interference

An shRNA previously demonstrated to knock down INF2 expression^3^ along with a control blank insert were cloned into a vector co-expressing farnesylated mApple (Addgene #54899). The shRNA and mApple were under control of the H1 and PGK promoters respectively. In-Fusion cloning was used to combine the regions of the original construct and the farnesyl-mApple into an AAV backbone. The following three sets of infusion primers were designed flanking the m-apple farnesyl and the shRNA and blank portions of the original plasmids:

#### shRNA

Forward: CTATACGAAGTTATGGTCGACGGTATCGATAAGCTTG; Reverse: GGTGGCGACCGGTAAGCTA.

#### m-Apple

Forward: TTACCGGTCGCCACCATGGTGAGCAAGGGC; Reverse: TTGATTCATAACTTCTCAGGAGAGCACACACTTG.

#### WPRE

Forward: GAAGTTATGAATCAACCTCTGGATTAC; Reverse: GTGGCAATGCCCCAA.

### *In vivo* cortical microinfarct induction

The animal’s head was affixed to a stable imaging apparatus under the two-photon microscope. The green laser light (532 nm) greatly under-fills the back aperture of the 20x objective (Olympus; XLUMPlanFI), yielding a fixed laser focus with ∼20–40 μm diameter at the imaging plane. The power of the green laser was ∼1 mW at the sample. We administered 25 μl of 5% w/v fluorescein-dextran (2 MDa) into the infraorbital vein over a period of 20 sec. The vasculature was clearly visible with two-photon imaging immediately after injection. The filter set used to detect fluorescein-dextran emission was 525/70m-2P (Chroma Corp). When finding a capillary location of interest, we targeted a region at least 20 μm away from larger penetrating vessels. After injecting 25–50 μl of 1.25% Rose Bengal into the infraorbital vein, using the same method described above for fluorescein-dextran we initiated photothrombosis by activating the green laser (1 mW at the sample) and allowing irradiation for 60-90 sec. Vessel occlusion was assessed by direct visual verification. In sham animals, green laser irradiation, at a typical power, led to no visible effects on the target vessel in the absence of rose Bengal.

### *In vivo* agarose controlled drug delivery

We generated acute, skull-removed cranial windows. Under 4% isoflurane anesthesia, we first injected 50 μL of 0.06 mg/mL buprenorphine (0.1 mg/kg for a 30 g mouse) intraperitoneally. Once the animal was in the surgical plane of anesthesia (4% MAC induction, 1-2% during surgery) the scalp was excised and periosteum cleaned from the skull surface. C&B MetaBond quick adhesive cement (Parkell; S380) was then applied to the skull surface to affix a custom-made metal flange to the right half of the skull. This metal flange could later be screwed into a custom holding post for head-fixation during imaging. A 3 mm diameter circular craniotomy (dura intact) was created over the left hemisphere, and centered over 1.5 mm posterior and 3 mm lateral to bregma, which encompasses the barrel and other regions of the somatosensory cortex. The cortical surface was cleaned of any blood and covered with a drop of warm 1.5% agarose dissolved in modified artificial cerebral spinal fluid ^4^, and immediately overlaid with a 4 mm diameter glass coverslip (Warner Instruments; 64-0724 (CS-4R). Care was taken not to compress the cortex during this process, which could affect cortical microvascular flow. MetaBond was then used to seal the edges of the coverslip, and to cover any remaining exposed skull surface or skin. We opted to use topical SMIFH2 application, rather than systemic injection. This was done to avoid confounding reductions in cerebral perfusion pressure that can occur with systemic dosing. SMIFH2 was dissolved in DMSO and then added to the warm 1.5% agarose solution (0.1% DMSO concentration) used in the cranial window procedure, providing direct access to the brain through the cortical surface. The final SMIFH2 concentrations in the agarose (150 μM) was chosen based on: (1) on the concentrations we tested on dissociated neurons in culture and (2) a study showing that ∼10% of similarly-sized molecules at the meninges enters the brain parenchyma through transcranial diffusion ^5^. Photothrombosis was initiated 30 min after starting the SMIFH2/vehicle loaded agarose application to allow diffusion through the cortex. Animals were perfused with PFA 4h post-stroke to harvest the brain.

### *In vitro* induction of actinification by bath applied NMDA

Three week old dissociated hippocampal cultures were incubated with either vehicle (H_2_O) or 50 µM NMDA added directly to their conditioned medium at 37°C for 5 min or other times, as indicated.

Neurons were kept in the culturing incubator prior to fixation. This approach was chosen over replacing conditioned media with fresh media containing NMDA due to the established toxicity of this manipulation ^6^. The following drugs were bath applied to the cultures at the following final concentrations: SMIFH2 75 µM, CK666 50 µM, APV 100 µM, MK 801 25 µM, C646 25 µM, Lat-A 2 µM, DIDS 500 µM, bumetanide 100 µM, GlyH-101 50 µM, NPPB 200 µM, JASP 4 µM.

### Ca^2+^ and Na^+^ depletion experiments

Conditioned medium was replaced either with calcium free solution in mM: HEPES 20, NaCl 137, MgSO_4_ 0.4, MgCl_2_ 0.5, KCl 5, KH_2_PO_4_ 0.4, NaH_2_PO_4_ 0.6, NaHCO_3_ 3, Glucose 5.6, EGTA 0.020, pH 7.3 (NaOH); or with sodium free solution: N-methyl-D-glucamine chloride (NMDGCl) 140.6, HEPES 20, MgSO_4_ 0.4, MgCl_2_ 0.5, KCl 1.4, KH_2_PO_4_ 1, KHCO_3_ 3, glucose 5.6, CaCl_2_ 1.8. pH 7.3, 320 mOSM. 50µM NMDA plus 10µM glycine were then added to induce actinification.

### Oxygen and Glucose deprivation

To induce oxygen and glucose deprivation 2-deoxyglucose was added to the cultures just before placing them into a hypoxic chamber XVIVO system (Biospherix, Parish, NY) at 37°C containing 1%O_2_.

### Neuronal cell culture transfection

Lipofectamine 2000 was used to transfect 3 weeks old dissociated hippocampal cultures, using various plasmids listed in the above key resources table.

Fresh neurobasal media was used to prepare the mixture of Lipofectamine and cDNA. 1 µl of Lipofectamine and 1.5 µg of cDNA per 50µl of NBM were mixed, incubated for 30 min and then added dropwise to 500 µl NBM in which neurons had been growing for 3 weeks. Washing was not required and neurons showed no overt signs of toxicity.

### Staining

#### Hippocampal cultures

Cultures were fixed with 3.7% formaldehyde in phosphate-buffered saline (PBS) plus 120 mM sucrose for 20 min at 37°C. Samples were incubated in 20 mM glycine for 5 min, rinsed and permeabilized with 0.2% Triton X-100 for 5 min at room temperature, and then blocked for 30 min with 2% bovine serum albumin (BSA). All primary antibodies were incubated for 1 hr at room temperature, and, following rinsing with PBS, were incubated with AlexaFluor-conjugated secondary antibodies (Invitrogen, Molecular Probes) for 45 min at 37°C. To label F-actin, AlexaFluor488-, 568- or 647-phalloidin at 1∶1000 (Invitrogen, Molecular Probes) was incubated for 2 hr at room temperature in the presence of 2% BSA. Finally, the coverslips were washed twice with PBS and mounted using either Aqua-Mount (Thermo Scientific) for acquisition with regular spinning disk confocal or Prolong Gold Antifade (Molecular Probes) for Airyscan acquisition.

#### Staining for cofilin-actin rods

Cultures were processed according to procedures modified from Mi et al, 2013^7^. Cultures were fixed for 45 min, at room temperature, in 4% formaldehyde in phosphate buffered saline (PBS) + 0.1% glutaraldehyde. Neurons were then permeabilized with −20°C cold methanol for 3 min, and then blocked with 5% goat serum/1% bovine serum albumin in TBS (10 mM Tris pH 8.0, 150 mM NaCl). Anti-actin and anti-cofilin antibodies were incubated overnight at 4°C, together with anti-MAP2. After rinsing with TBS, neurons were incubated with AlexaFluor-conjugated secondary antibodies (Invitrogen, Molecular Probes) for 1h at RT. Finally, the coverslips were washed again with TBS and mounted using ProLong Gold Antifade (Molecular Probes).

#### Brain section histology and analyses

For immunohistochemical analysis, animals were perfused transcardially with 4% paraformaldehyde in phosphate-buffered saline (PBS), pH 7.4. The brains were then placed in 30% sucrose overnight and shipped to the Halpain lab where the brain region caudal to the bregma was sliced into 30 µm-thick coronal sections by using a vibratome, and stored in PBS. After blocking for 2 hours at room temperature with 20% normal goat serum (NGS) and 0.3% TritonX-100 in PBS, sections were incubated with primary antibodies in 10% NGS and 0.3% TritonX-100 for 24h at 4°C on an orbital shaker. Afterwards the sections were rinsed in PBS three times over a period of three hours, before being incubated with the secondary antibodies overnight at 4°C. Finally, the floating sections were transferred from the solution with the secondary antibodies to a solution with phalloidin and DAPI for the last 24h before washing and mounting them with Aqua-Mount (Lerner Laboratories) on positive charged slides.

#### Platinum Replica Electron Microscopy

Sample preparation for platinum replica electron microscopy was performed as described previously ^8, 9^. In brief, detergent-extracted samples were sequentially fixed with 2% glutaraldehyde in 0.1 M Na-cacodylate buffer (pH 7.3), 0.1% tannic acid, and 0.2% uranyl acetate; critical point dried; coated with platinum and carbon; and transferred onto 50 mesh electron microscopic grids for observation. Detergent extraction was done with 1% Triton X-100 in PEM buffer (100 mM Pipes-KOH, pH 6.9, 1 mM MgCl_2_, and 1 mM EGTA) containing 2% polyethelene glycol (molecular weight of 35,000), 2 μM phalloidin, and 10 μM taxol for 3 min at room temperature. Samples were analyzed using JEM 1011 transmission electron microscope (JEOL USA, Peabody, MA) operated at 100 kV. Images were captured by ORIUS 832.10W CCD camera (Gatan, Warrendale, PA). PREM images are presented in inverted contrast. Color labeling was performed using Hue/Saturation tool in Adobe Photoshop to avoid obscuring the structural details.

### Image acquisition

Unless otherwise indicated, all images shown in this article represent maximum projection images derived from a z-stack.

### Fixed specimens: Confocal spinning disk

To acquire images of fixed dissociated cultures or brain sections we used an Olympus IX-70 microscope equipped with a CSU-X1 spinning disk confocal (Yokogawa Electric Corporation) custom equipped with 405 nm, 491 nm, 561 nm and 640 nm 50 mW solid state lasers (Solamere Technology Group Inc.) and a CoolSNAP HQ2 digital CCD camera (Photometrics) with pixel size of 91 nm. Fluorescence emission was selected through the following bandpass filters: 525/50 nm, 595/50, 700/75. Metamorph (Molecular Devices) was used to acquire a stack of 6-11 images in the z-dimension using optical slice thickness of 0.2 µm for 60X images of dissociated cultures and brain sections. A single plane of focus was used to acquire low magnification images of brain sections using a 1.25X objective.

### Fixed specimens: Super-resolution imaging

A Zeiss LSM 880 Rear Port Laser Scanning Confocal with Airyscan FAST Microscope with a 63X/1.4 oil objective was used to acquire super-resolution images of actinified neurons (Fig 2A).

### Time-lapse imaging

Live images were acquired every 30 s before and after NMDA-induced actinification or every 15 minutes when recovery from actinification was monitored with image acquisition times of 0.01–0.2 s using a Nikon Ti-E microscope with perfect focus system (Nikon) and an iXon X3 DU897 EM-CCD camera (Andor Technology plc). The microscope was equipped with a CSU-X1 spinning disk confocal (Yokogawa Electric Corporation), and a customized CO_2_-delivery, temperature-controlled chamber (5% CO_2_, 35°C). A 60X 1.4 NA Plan APO oil immersion objective was used for all experiments. Fluorescent specimens were excited using a laser launch (Solamere Technology Group Inc.) equipped with 488 nm, 561 nm and 640 nm 100 mW solid state lasers. Fluorescence emission was selected through the following band-pass filters: 525/50 nm, 595/50, 700/75. Metamorph (Molecular Devices) was used to acquire a stack of images in the z dimension using optical slice thickness of 0.2 µm; during live imaging the number of z-sections was typically kept to ≤4 to minimize photodamage. Only experiments in which focus was precisely maintained throughout the recording session were included in the time-lapse analyses. Experiments that required imaging neurons for long durations during cell stress, such as monitoring recovery after actinification, required neurons to be cultured and transfected on 35 mm glass bottom petri-dishes (Matsunami) for use with our Tokai HIT CO_2_-delivery, temperature-controlled chamber (5% CO_2_, 35°C), with an embedded heated water bath to tightly control humidity and temperature. Movies were acquired on a Nikon system equipped with CSU-W1 spinning disk, photometrics prime 95B sCMOS camera and Nikon Live SR optical super resolution module.

### Cell viability assays

#### VivaFix staining in dissociated cultured neurons

VivaFix reagent was added directly to the conditioned media and cultured neurons for 20 min at 37°C, modifying manufacturer’s protocol (Biorad). Samples were rinsed once with conditioned media from sister cultures before live imaging or fixing the cells, and co-staining with phalloidin or other markers where indicated.

#### Fluoro-Jade C staining in brain sections

Free-floating brain sections were first incubated with the anti NeuN antibody for 24 h. Then they were mounted onto charged slides, which were dried for 30 min at 50°C, then rinsed for 5 min in distilled water, incubated in 0.06% potassium permanganate solution for 5 min and rinsed again in water. Slides were then transferred for 10 min to a 0.0001% solution of Fluoro-Jade C dissolved in 0.1% acetic acid. DAPI was added at this step. The slides were then rinsed through three changes of distilled water for 1 min per change. Excess water was drained, and slides were then air dried on a slide warmer at 50°C for at least 5 min. The air-dried slides were cleared in xylene for at least 1 min and then coverslipped with DPX non-fluorescent mounting media.

### Quantification

#### Image analysis

In all experiments digital images were acquired using identical parameters and settings (e.g., laser excitation power, acquisition time, time-lapse interval, exposure time, etc.) across experimental conditions. All images displayed in this paper use identical image display settings whenever experimental groups are compared to one another. Sample sizes are provided in all figure legends. For analyses we used Fiji, the open source image software ^10^. Adobe Systems Inc. software Photoshop was used for image display, and for analysis of fixed cultures using the binary scoring system described below.

#### Quantification of actin reorganization (“actinification”) in cultured neurons

To quantify the degree of neuronal F-actin reorganization under various conditions, we devised a convenient assay based on the percentage of neurons that exhibited altered F-actin distribution. We used a binary scoring system that identified individual neurons as being either “actinified” or “non-actinified,” as visualized in fixed cultures using fluorescent phalloidin as a label for F-actin. “Actinified” neurons were defined as those that showed robust accumulation of filamentous-appearing phalloidin staining within the soma and proximal dendrites, with little or no phalloidin accumulation in a spine-like punctate pattern; “non-actinified” neurons showed robust accumulation of phalloidin staining in dendritic spine protrusions, and only faint, non-filamentous phalloidin staining in the soma and dendrites. This binary mode of designation was valid because, although different individual neurons became actinified at various times following the onset of a stimulus (e.g., NMDA), once actinification was initiated it proceeded rapidly, and the filaments that accumulated in actinified neurons were stable for long periods. This subjective, binary approach to quantification was applied in an unbiased manner -- observers assigned to collect and quantify the images were blind to the experimental manipulation via randomized encoding of the samples. Sample identification was not unveiled until after all data for a given experiment were collected and analyzed.

In Fig 3B and C actinification was quantified within the proximal region of the dendrite outlined by Fiji. Briefly, the proximal region of the shaft was cropped after subtracting background. The cropped region of the shaft was then thresholded so that it would be uniformly highlighted. The thresholded area was then binarized and any feature outside the shaft was removed using the erase tool. The plugin Macros Macros was used to outline the binarized thresholded area to generate a ROI (region of interest), which could be saved and added to the original cropped image using ROI Manager, as shown in Fig 3C. This allowed Fiji to measure intensity only within the generated ROI. In Fig 4A, dendritic actinification was quantified using live cell imaging of individual neurons labeled using the genetically encoded fluorophores mApple-F-tractin, Lifeact-mRFP, or SiR-Actin, which selectively label F-actin. Using time-lapse images collected over the course of 20 to 30 minutes, a region of interest (ROI), small enough to fit between spines perpendicular to the plane of focus of the shaft, was used to quantify, over time, the F-actin signal intensity within the background-corrected dendrite shaft, which increased during actinification, as quantified over time by a line scan (1 pixel wide) manually drawn across the same dendrite labeled with the co-transfected plasma membrane marker (lck-GFP). In Fig 2C we used the Nikon software NIS Elements and F-actin intensity was measured inside small ROIs placed both in the dendritic shaft and inside adjacent spines within an image. The dimensions of the ROIs were determined by the ability to remain within the individual morphing spines over time or as far as the shaft, by the presence of spines perpendicular to the plane of focus.

### Brain sections

Actinified neurons were identified within the YFP (+) neuronal population by observers blinded to the specific experimental conditions, but trained to recognize the characteristic somatic organization of the actin cytoskeleton. A neuron was defined as “actinified” if it displayed filamentous-like phalloidin staining in the interior of the soma and proximal dendrite, and “non-actinified” if it lacked clear signs of such a pattern. The quantification was organized by regions of the cortex, based on their relative proximity to the ischemic core. The core of the infarct was defined as the region with strongly reduced staining of NeuN (< 90% staining intensity relative to control); the penumbra region was defined as the zone showing robust NeuN staining intensity and radially surrounding the core within 170-250 µm.

FluoroJade (+) cells in selected regions of the cortex were quantified using the Fiji “analyze particles” function to automatically identify both the overall number of nuclei (DAPI) and the number of FluoroJade and DAPI (+) cells, with values corrected for area To quantify the effect of the formin inhibitor SMIFH2 versus vehicle on formation of cavitites within the infarct, we calculated the ratio of the area of the cavities (regions lacking neuropil) to the area of ischemic damage, defined as the region with robustly depleted NeuN immunostaining (the “core” of the infarct, as described above). We also quantified the area of vascular leakage as the region staining for mouse immunoglobulin G. The area of tissue damage was measured in adjacent tissue sections and the total infarct volume, Vt, was calculated by Vt = (A_1_ + A_2_ + …+ A_n_) h, where A_n_ was the area of damage in the nth slice, and h was the distance between adjacent sections.

## Statistical Analyses

All results reported here were observed reproducibly in at least two to three independent culture preparations; similarly, stroke experiments *in vivo* were repeated across multiple days using mice from multiple litters. Prior to quantitative analysis, sample identity was either encoded for blinding of the experimental group prior to analysis, or image acquisition and analysis were conducted by different people to avoid observer bias. Statistical significance was set at the 95% confidence level (two tailed) and calculated using Prism (Graphpad Software). Values are presented as the mean ± S.E.M.

To assess whether the data were normally distributed we used the D-Agostino-Pearson test to determine deviation by skewness or kurtosis. When normality was not met an appropriate non-parametric test was chosen, as described in the legend of the corresponding figure. Adjustments for multiple comparisons were made by using one-way ANOVA, two-way ANOVA, and appropriate multiple comparison post-hoc tests, as stated in the corresponding figure legends. Pharmacological and genetic experiments were statistically compared to their corresponding vehicle or wild type control constructs.

## Supplemental Information

Supplemental information includes 8 figures and can be found with this article

## Supplemental Figure Legends

**Supplemental Figure S1.**
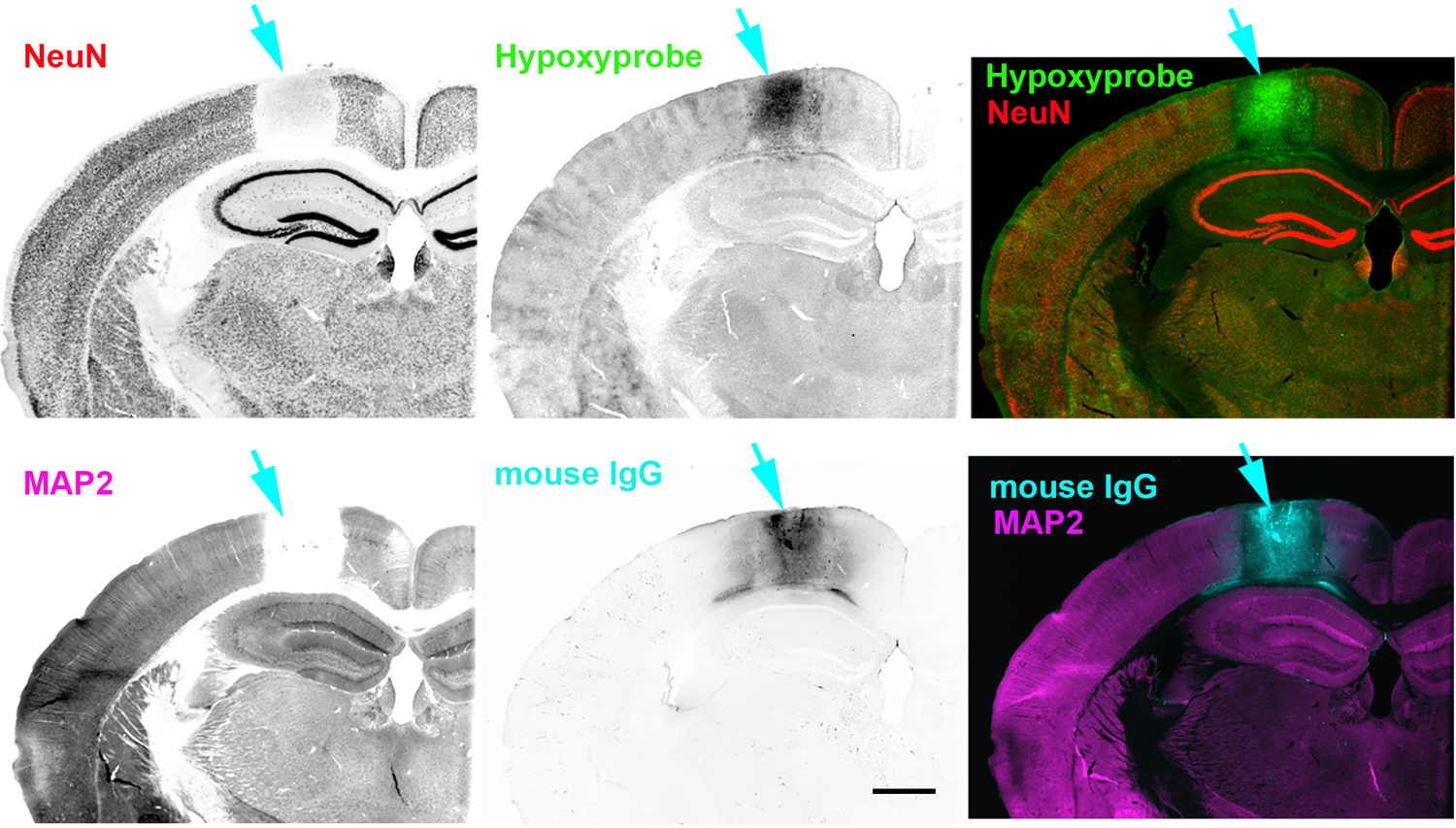
Selected histological markers of ischemic brain injury. Representative images of a, individual section of a mouse brain subject to single vessel photothrombotic stroke, immunostained with some of the indicated markers used in this study to characterize ischemic changes: anti-NeuN (specifically identifies neuronal cell bodies); anti-MAP2 (specifically identifies neuronal dendrites); hypoxyprobe (which uses an antibody to detect the presence of pimonidazole, which accumulates in tissues where pO2 < 10mmHG); and mouse anti-IgG (to assess the extent of hypoxia-induced vascular leakage and/or blood-brain-barrier disruption). Scale bar, 700 µm.

**Supplemental Figure S2.**
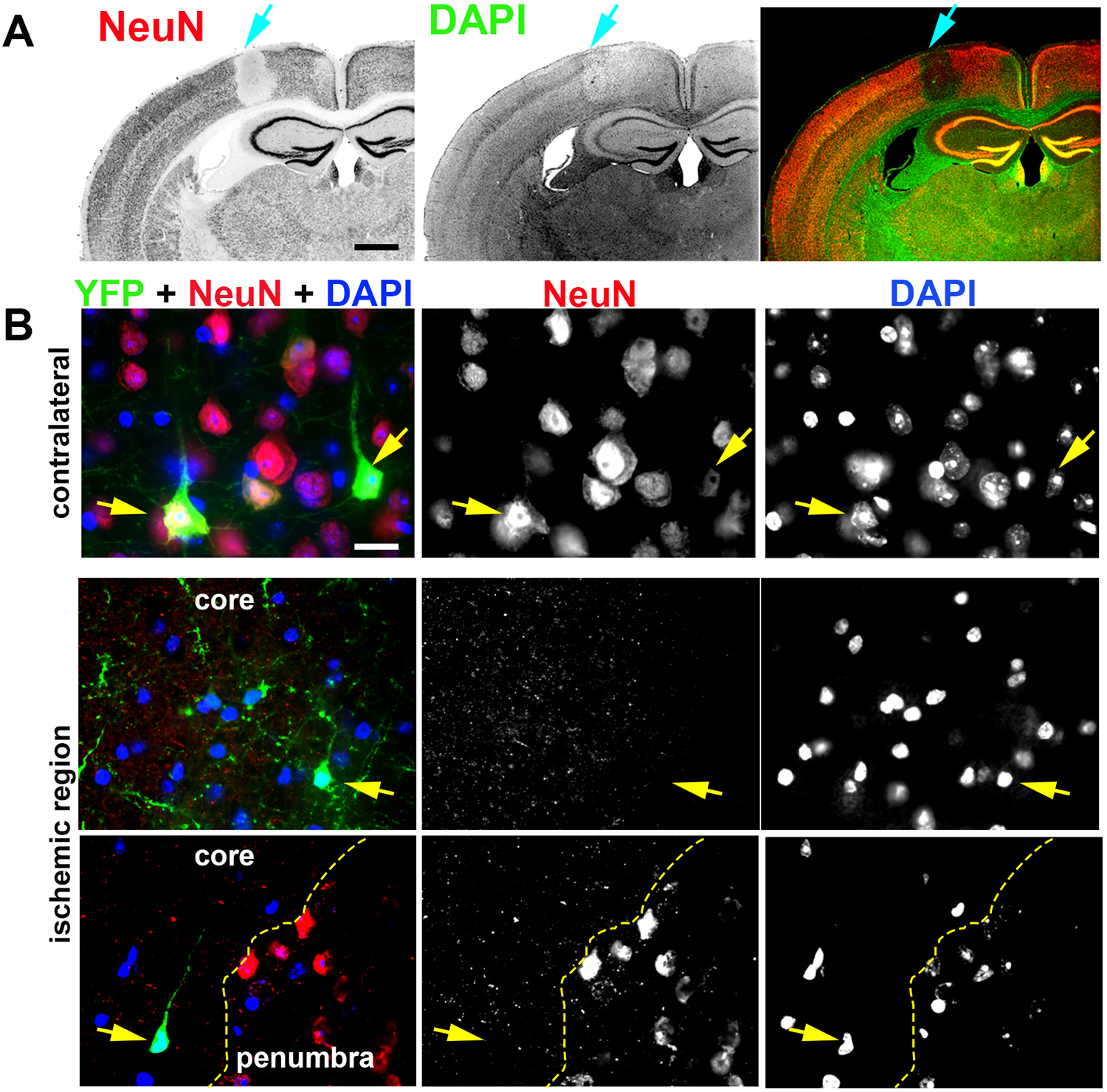
Reduction of NeuN immunofluorescence in the ischemic region does not reflect post-stroke neuronal loss. (**A**) Representative Thy1-GFP mouse brain section stained for NeuN and DAPI, 6h post-stroke, displaying a clear loss of NeuN signal in the ischemic core (indicated by the *blue arrow*), but relatively little reduction in DAPI staining. Scale bar, 1000 µm. (**B**) Selected regions of cortical layer 5/6 of somatosensory cortex (from the region indicated by the *blue arrow* in *A*) comparing at high magnification brain tissue ipsilateral vs. contralateral to the stroke. In the control, contralateral region (*top row*) DAPI and NeuN signals are strong and present together tin all neuronal cell bodies. *Arrows* indicate YFP-positive neurons. In contrast, within the ischemic core (*middle row*), NeuN staining is lost from individual neurons within the ischemic core, while DAPI staining persists. Thy1-driven soluble GFP is detectable in neurons both within and outside the ischemic core. *Dashed yellow line* in the bottom row indicates the boundary where NeuN staining is lost in the ischemic core, relative to staining in the surrounding presumptive penumbra, which can be seen at this magnification as a depletion of NeuN immunoreactivity in individual neuronal cell bodies within the core, but not the penumbra. Scale bar, 30 µm

**Supplemental Figure S3.**
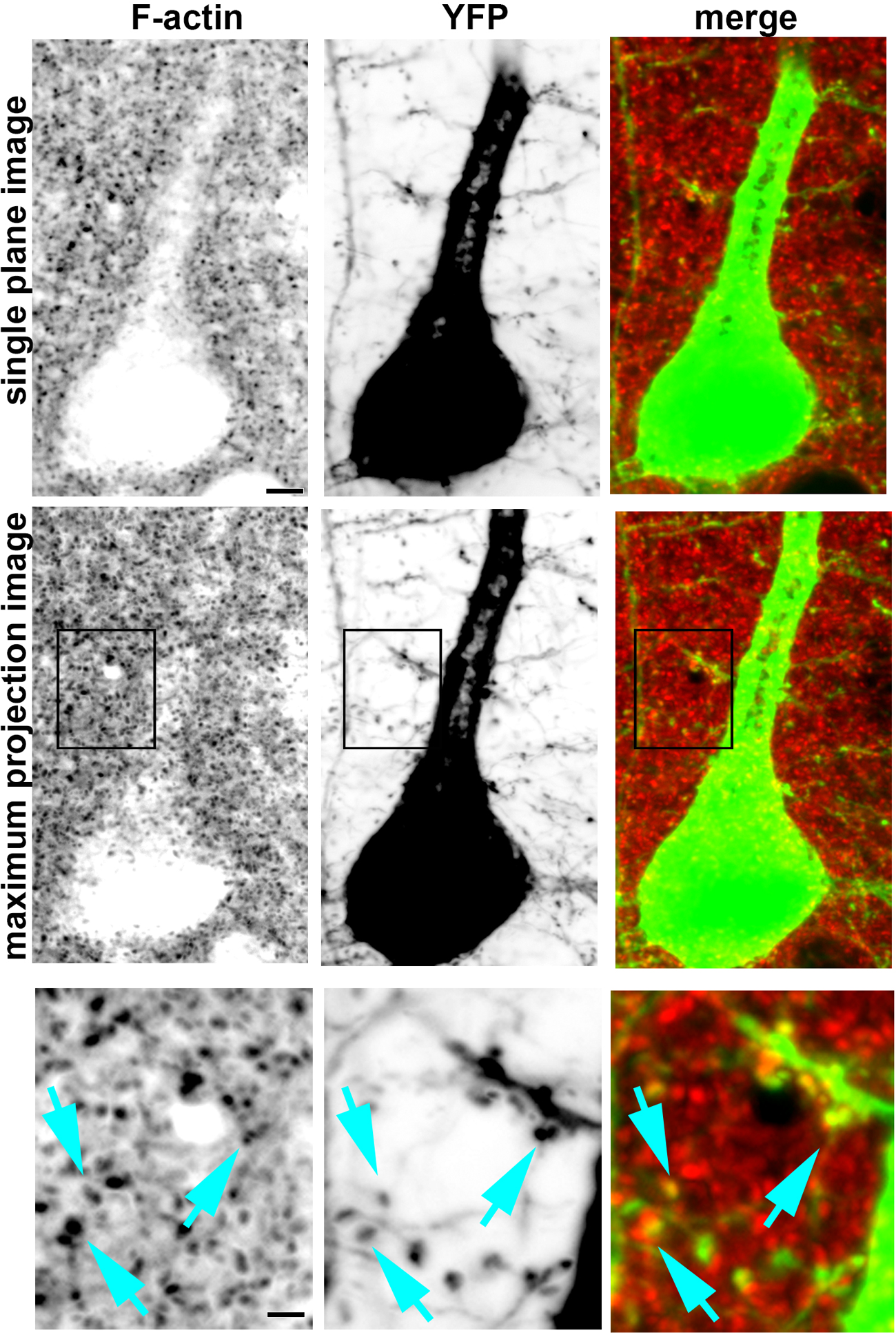
Enrichment of F-actin within dendritic spines in cortex of control mouse brain. A Thy1-YFP mouse brain was processed and stained for F-actin as described, using Alexa Fluor 647-phalloidin. Here we focus on a single YFP-positive pyramidal neuron and surrounding neuropil from Layer 5/6 in somatosensory cortex, imaged by confocal microscopy in coronal brain sections using a 60x (N.A. 1.42) oil immersion objective. Images in the *left column* show phalloidin staining in reverse grayscale; images in the *center column* show the YFP fluorescence in reverse grayscale; the *right column* shows a colorized merged image of the two. The *top row* of images show a single plane from the z-stack of collected images. Note that dense, punctate phalloidin staining is present throughout the neuropil, but the neuronal cell body and proximal apical dendrite are relatively devoid of phalloidin staining. The *middle row* of images shows the same x-y field of view but as a maximum projection image from the z-stack. Note the marked increase in the number and density of phalloidin puncta, and the fact that many such puncta now overlay portions of the soma and apical dendrite. We interpret this pattern to indicate that phalloidin-positive puncta densely surround the dendrite in three dimensions, but that the interior of the somatodendritic compartment itself is relatively non-enriched for F-actin under control conditions. (We elsewhere show that stroke substantially alters this pattern, as F-actin accumulates within the somatodendrite interior; see Figure 1). The *bottom row* shows the boxed region at higher magnification, illustrating our consistent observation that all dendritic spines colocalize with phalloidin-positive puncta; *cyan arrows* point to specific examples. Given the high density of dendritic spines within mammalian cortical neuropil, we conclude that the majority of phalloidin puncta in the size range of ∼0.2-1 µm in diameter that we observe are likely to correspond to dendritic spines, where F-actin is enriched. Scale bar, 4 µm *upper & middle rows*; 2 µm, *bottom row*.

**Supplemental Figure S4.**
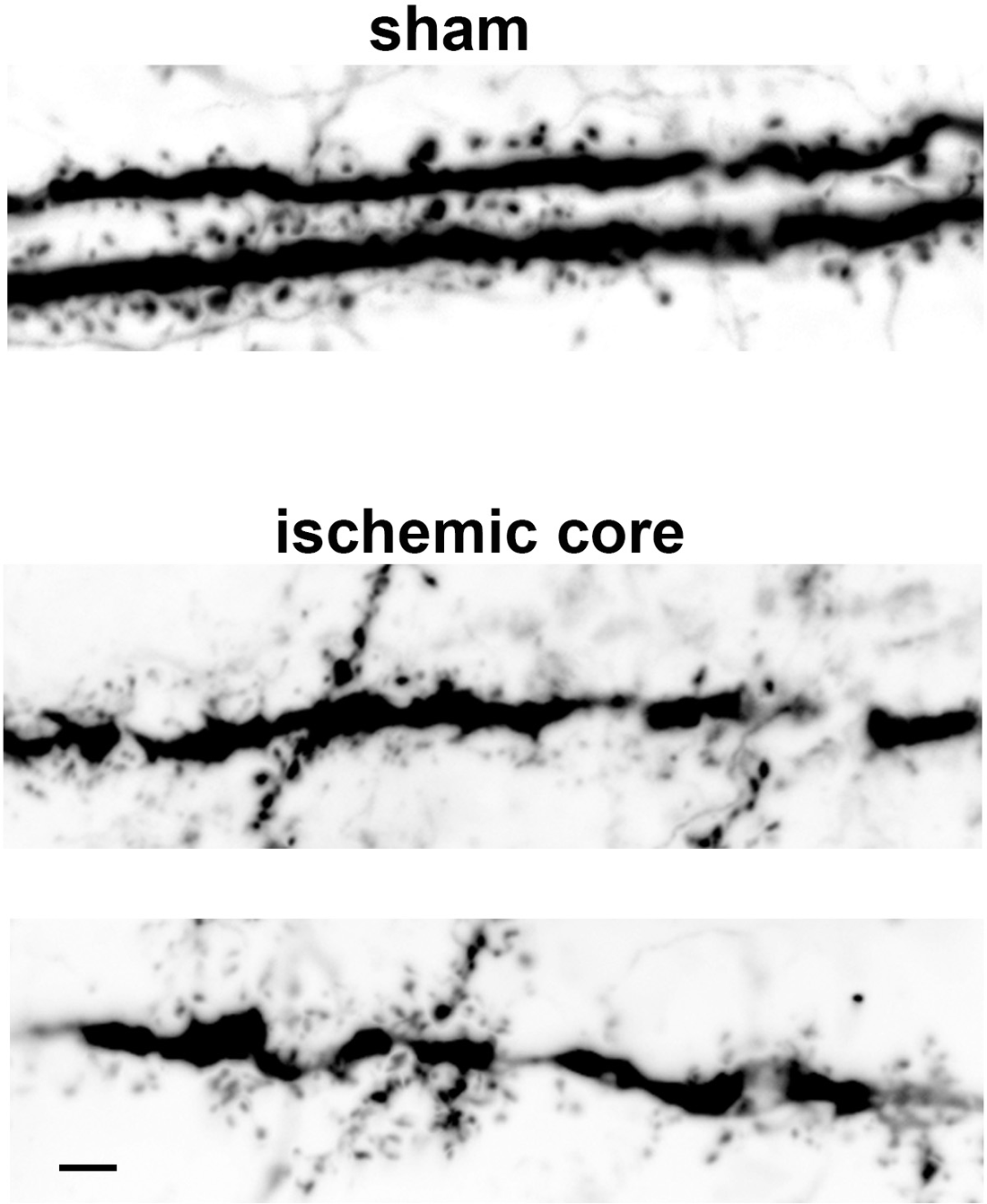
Dendritic spines decrease in size and number after stroke. Selected dendritic regions proximal to the cell bodies of Thy1-YFP-positive layer 2/3 pyramidal neurons from sham-operated vs. ischemic mouse brain. YFP fluorescence is displayed in reverse grayscale. Scale bar, 5 µm.

**Supplemental Figure S5.**
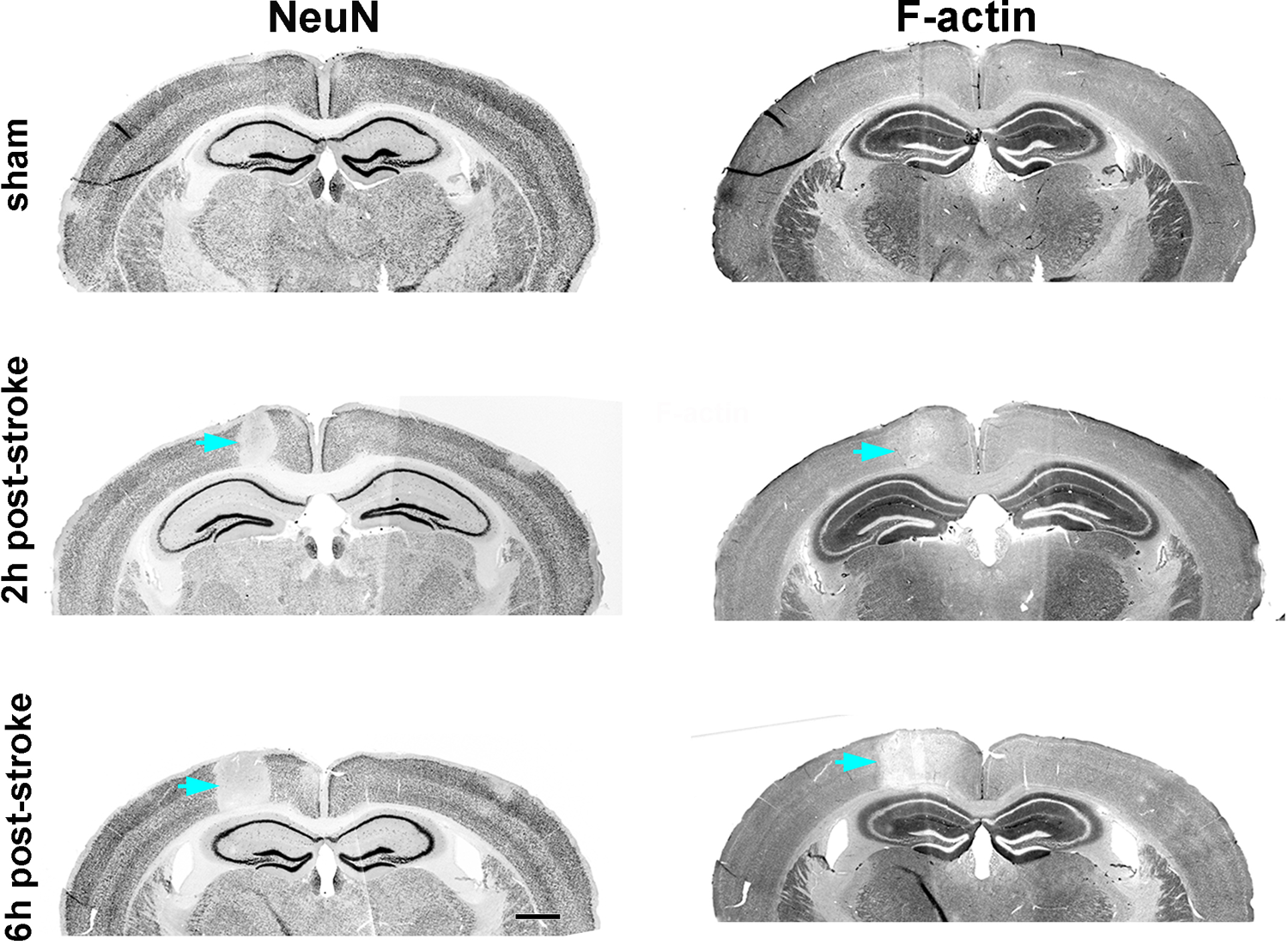
Overall F-actin signal within the ischemic core decreases over time following stroke. Representative tissue sections from sham or ischemic Thy1-YFP brains fixed at 2h and 6h after single vessel photothrombotic stroke and co-stained for NeuN and F-actin. *Cyan arrows* indicate the ischemic region. Note that a dramatic decrease in NeuN immunoreactivity is already observed by 2h after stroke, accompanied by a very modest decrease in phalloidin staining. By 6h post-stroke we observe that, as expected, the average infarct volume (as measured using NeuN or other markers) increases, and it is accompanied by a consistent decrease in overall phalloidin staining within the infarct zone. Nevertheless, within the infarct zone (core and penumbra) we observe numerous neuronal cell bodies with accumulations of somatodendritic F-actin that are suggestive of actinification (see Fig 1). Scale bar, 700 µm.

**Supplemental Figure S6.**
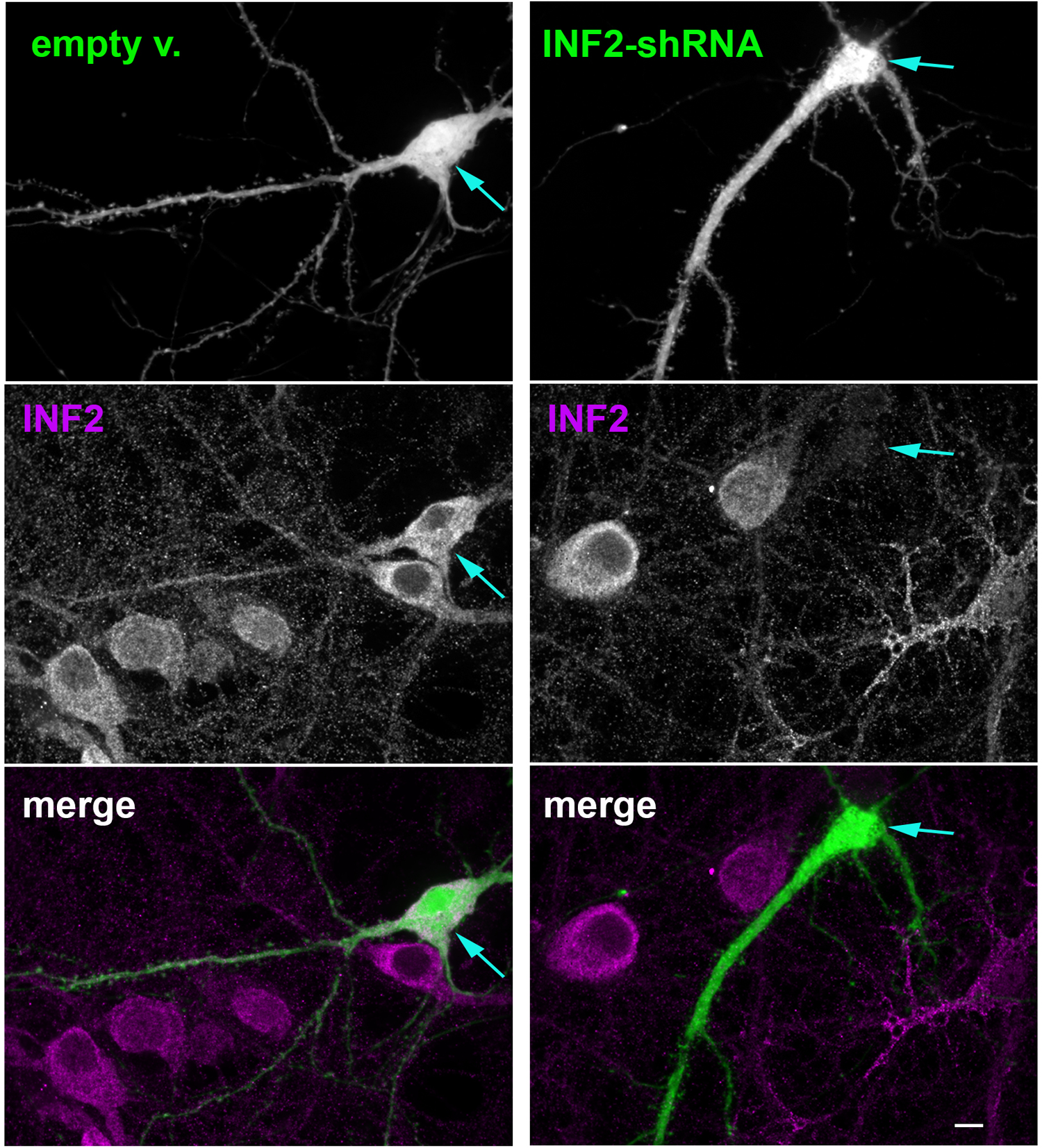
Specificity of INF2 immunoreactivity determined by RNA interference. Selected cultured hippocampal neurons transfected at DIV14 with either the empty vector (left) or the INF2-shRNA (right) for 10 days prior to fixation and immunostaining against endogenous INF2. *Blue arrows* point to the cell bodies of the transfected neurons and to the presence or lack of INF2 immunoreactivity in the two different experimental conditions. Scale bar, 8 µm.

**Supplemental Figure S7.**
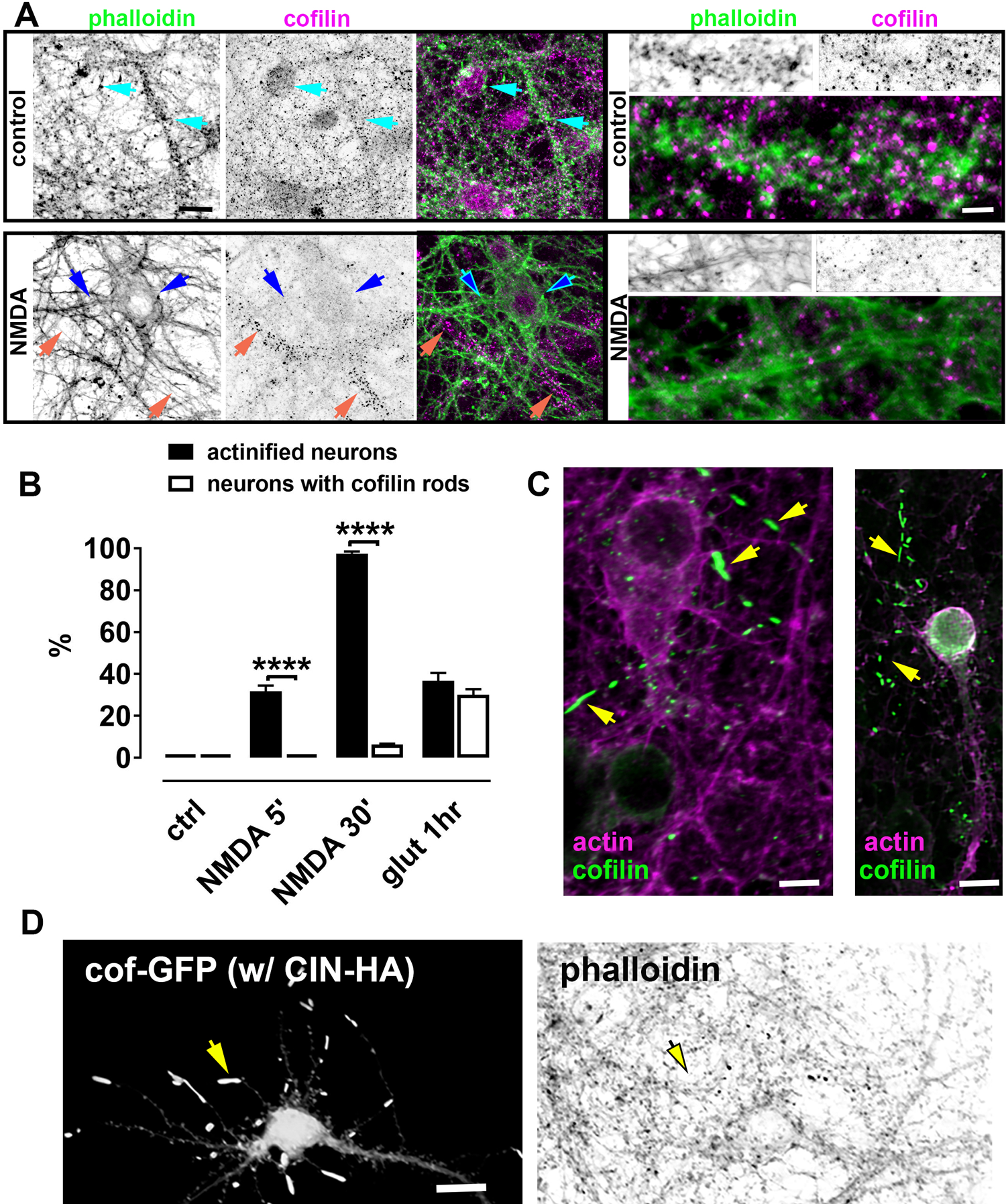
Somatodendritic actinification is a novel process distinct from cofilin-actin rod formation. (A) *Left*, representative black and white images of control and NMDA-treated neurons co-stained for F-actin, and cofilin. Endogenous cofilin shows the expected punctate distribution in control neurons (*cyan arrows*), and little detectable immunoreactivity in NMDA-treated actinified neurons (*blue arrows*), while remaining detectable and punctate in a neighboring astrocytes (*orange arrows*). Scale bar, 20 µm. *Right*, selected dendritic regions from a control neuron and an actinified NMDA-treated neuron. Scale bar, 4 µm. **(B)** Cofilin rod formation does not occur in parallel with actinification. Cofilin rods were not detected after incubation for 5 min with NMDA, and only rarely after 30 min of NMDA, even in the actinified neurons; in contrast, we reliably detect cofilin rods after incubation with 200 µM glutamate for 1 hr or longer. Data are mean ± SEM from three independent culture preparations; **** p< 0.0001; two-way ANOVA, followed by Sidak’s multiple comparisons *post-hoc* test, to selectively compare neurons with cofilin rods and actinified neurons within each experimental group. **(C)** Two representative images demonstrating the lack of colocalization between the actinified somatodendritic regions (endogenous actin stain detected using an anti-actin antibody (*magenta*); endogenous cofilin rods, detected using an anti-cofilin antibody (*green*). Scale bar, 10 µm (*left* image); 18 µm (*right* image). **(D)** Ectopic co-expression of cofilin and chronophin, one of its Ser-3 phosphatases, induces cofilin rods without inducing actinification. Representative neuron transfected with cofilin-GFP (Cof) and chronophin-HA (CIN) displaying the distribution of cofilin rods in distal regions of thin secondary dendrites (*arrows*). Scale bar, 24 µm.

**Supplemental Figure S8.**
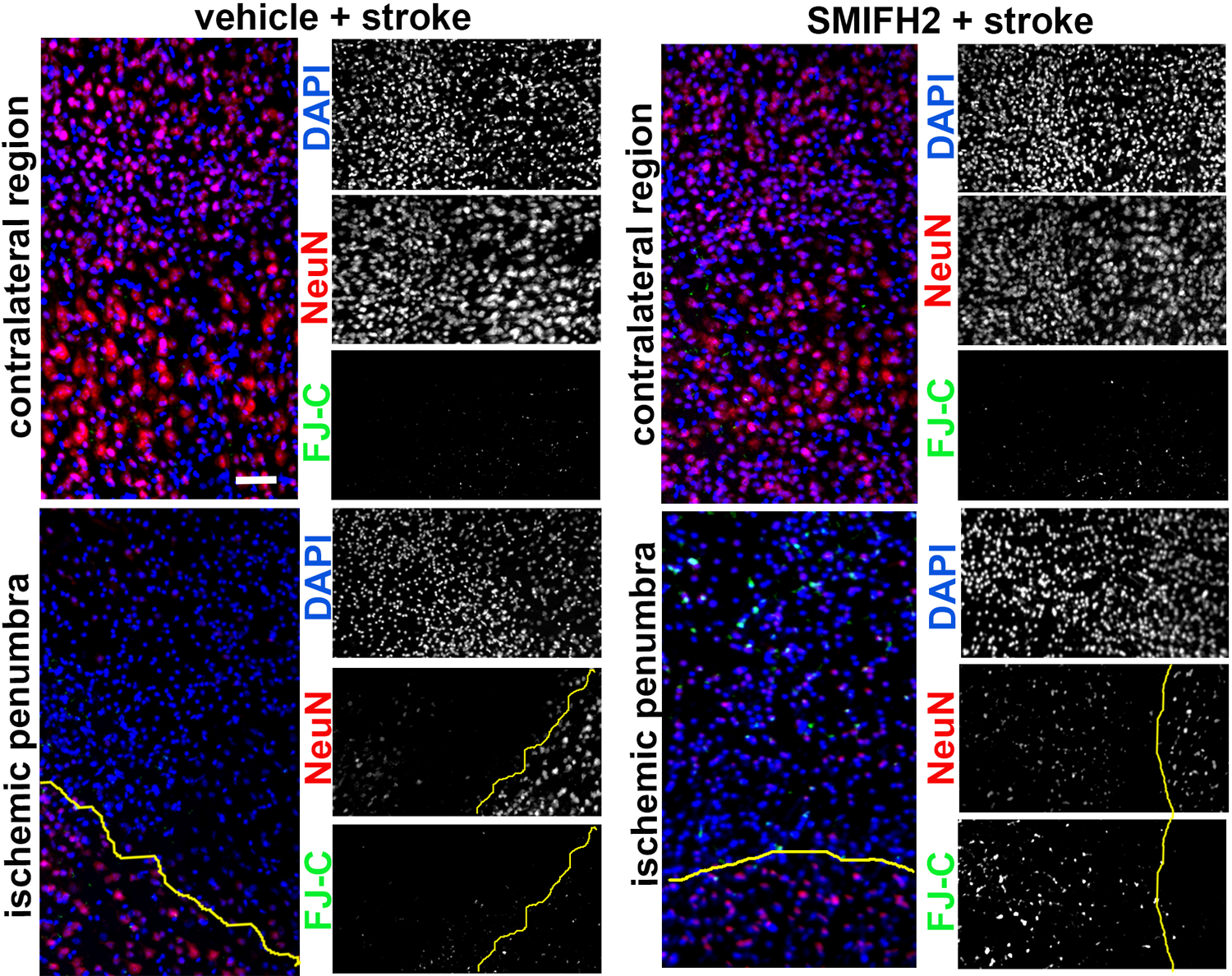
Formin inhibition increases early cell death in the ischemic penumbra. Representative cortical regions from the hemipshere contralateral to the infarct (*upper half*), and along the border (*yellow line*) between the infarct core and the penumbra (*lower half*), in the absence (*left*) or presence (*right*) of formin inhibitor SMIFH2 delivered by passive diffusion from a saturated agar plug placed over the thinned skull. Each color-combined image is juxtaposed to the correspondent individual grayscale channels (which are rotated 90 deg relative to the merged image). Nuclei are labeled with DAPI (*in blue*), NeuN (*in red*), and the cell death marker FluoroJade-C (FJ-C; *green*). Scale bar, 60 µm.

## Notes

### Competing Interest Statement

The authors have declared no competing interest.

